# Epithelial cell chirality emerges through the dynamic concentric pattern of actomyosin cytoskeleton

**DOI:** 10.1101/2023.08.16.553476

**Authors:** Takaki Yamamoto, Tomoki Ishibashi, Yuko Mimori-Kiyosue, Sylvain Hiver, Naoko Tokushige, Mitsusuke Tarama, Masatoshi Takeichi, Tatsuo Shibata

**Author notes:** These two authors contributed equally (Takaki Yamamoto, Tomoki Ishibashi).

## Abstract

The chirality of tissues and organs is essential for their proper function and development. Tissue-level chirality derives from the chirality of individual cells that comprise the tissue, and cellular chirality is considered to emerge through the organization of chiral molecules within the cell. However, the principle of how molecular chirality leads to cellular chirality remains unresolved. To address this fundamental question, we experimentally studied the chiral behaviors of isolated epithelial cells derived from a carcinoma line and developed a theoretical understanding of how their behaviors arise from molecular-level chirality. We first found that the nucleus rotates and the cytoplasm circulates robustly in a clockwise direction. During the rotation, actin and myosin IIA are organized into stress fibers with a vortex-like chiral orientation at the ventral side of the cell periphery, simultaneously forming thin filaments with a concentric orientation at the dorsal level of the cell. Surprisingly, we found that the intracellular rotation is driven by the concentric pattern of actomyosin filaments on the dorsal surface of the cell, not by the vortex-like chiral stress fibers. To elucidate how the concentric actomyosin filaments induce chiral rotation, we analyzed a theoretical model developed based on the theory of active chiral fluid, and revealed that the observed cell-scale unidirectional rotation is driven by the molecular-scale chirality of actomyosin filaments even in the absence of cell-scale chiral orientational order. Our study thus provides novel mechanistic insights into how the molecular chirality is organized into the cellular chirality and an important step towards understanding left-right symmetry breaking in tissues and organs.

## INTRODUCTION

Left-right asymmetry is ubiquitously observed in the bodies and organs of organisms. Despite extensive research, however, we still do not have a complete understanding of how left-right asymmetric structures are formed at an organismal scale. The breaking of left-right symmetry at the body and organ scale has been investigated in embryonic bodies, such as early vertebrate embryo [1, 2] nematodes [3–5] and pond snails [6–8]; and in organogenesis, such as embryonic hindgut [9–11] and male genitalia [12] in *Drosophila* and heart-looping in the chicken [13]. Interestingly, in most of these cases, the left-right symmetry breaking at the organ scale is associated with chiral features at the cellular scale, indicating that cell-level chirality induces multicellular chirality [14]. Chiral dynamics have been observed in isolated single cells, such as nerve cells [15], zebrafish melanophores [16], human foreskin fibroblasts (HFF) [17, 18], and Madin-Darby canine kidney (MDCK) cells [19]. Furthermore, experimental and theoretical studies have revealed that cell-intrinsic chirality drives left-right asymmetric morphogenesis of tissues [20–22] and organs [13]. Therefore, to elucidate the mechanism of left-right symmetry breaking of organismal structures, it is essential to comprehensively investigate the mechanism underlying chiral dynamics at the single-cell scale.

In cells, there are many chiral components, such as amino acids, proteins, and DNA, and their proper organization can induce chiral properties of cells [23]. Particularly, cytoskeletal molecules such as actin and microtubules have been suggested as candidate apparatuses driving chiral dynamics at the single-cell scale. For instance, actin and myosin are responsible for the chiral nuclear rotation of zebrafish melanophore [16], and the chiral neurite extension in nerve cells [15]. Actin polymerization by formin drives chiral swirling of the cytoskeleton in HFF [17]. Microtubules are involved in the chirality of neutrophils [24]. These studies attribute the chiral cell dynamics to the chiral rotating dynamics of actin and microtubules driven by molecular motors [25, 26]. However, a mechanistic understanding of how these molecules generate cell-scale chirality is still not complete. Several attempts to gain mechanistic insights using theoretical models indicate that the chiral symmetry breaking at the cellular level requires spatial coordination of chiral cytoskeletal molecules [3, 17]. In particular, for the chirality of HFF, it has been proposed that the transverse actin fibers physically interact with radial actin fibers, which are screwed by formin, to drive the nuclear rotation in the counterclockwise direction [17, 18]. In *C. elegans* embryo, it has been proposed that the actomyosin cortex generates active chiral torque and its spatial gradient induces the chiral symmetry breaking [3]. Therefore, to crack the code of cellular chirality, it is important to elucidate how molecular-scale chiral activity spatially coordinates to trigger cellular-scale chirality.

In the present study, we investigated the behavior of Caco2 cells, a typical epithelial cell line that was derived from colorectal adenocarcinoma. We found that, when these cells were singly isolated and cultured on substrates, the nucleus rotates along with the circulation of the cytoplasm in a clockwise direction, as viewed from above. We then showed that actin and myosin II are responsible for this rotation of intracellular components. These cytoskeletal molecules formed concentric actomyosin filaments at the dorsal side of the cells, while they were organized into stress fibers with a vortex-like chiral orientation at the ventral side. Our experiments suggest that the former structure most likely drives the rotating motion, implying that a cell can rotate without any cell-scale chiral orientational order of the cytoskeleton. To elucidate whether the concentric achiral pattern of the actomyosin filaments can indeed generate rotational flow, we analyzed a hydrodynamic model, based on the active chiral fluid theory, of a three-dimensional (3D) cell, considering the effect of molecular chirality of actin and myosin. We found that the concentric achiral structure of actomyosin can generate chiral cytoplasmic circulation, due to the force which originates from the molecular chirality of individual cytoskeletal components, even without cell-scale chiral structures. On the other hand, we found no evidence that radial actin fibers are involved in the nuclear rotation in Caco2 cells, suggesting that there might be cell type-specific mechanisms to rotate cytoplasmic components.

## RESULTS

### Nuclei of singly isolated Caco2 cells rotate in a clockwise direction

To study the rotational dynamics of epithelial cells, we cultured singly isolated Caco2 cells and imaged them using a differential interference contrast (DIC) microscope (Fig. 1A, Movie S1). 76% of isolated Caco2 cells spread circularly on a collagen-coated glass substrate, generating lamellipodia in all directions along the cell periphery with no persistent migration [27]. In the cells spreading circularly, we noticed that the nuclei exhibit rotational motion in a clockwise direction when viewed from the dorsal (apical) side (Fig. 1B). There was no cell that exhibited rotational motion in a counterclockwise direction. 24% of the cells exhibited migratory behavior at the start of our live imaging, and it took a while for the cells to spread circularly without persistent migration. The cells exhibiting migratory behavior were excluded from the analysis.

**FIG. 1.**
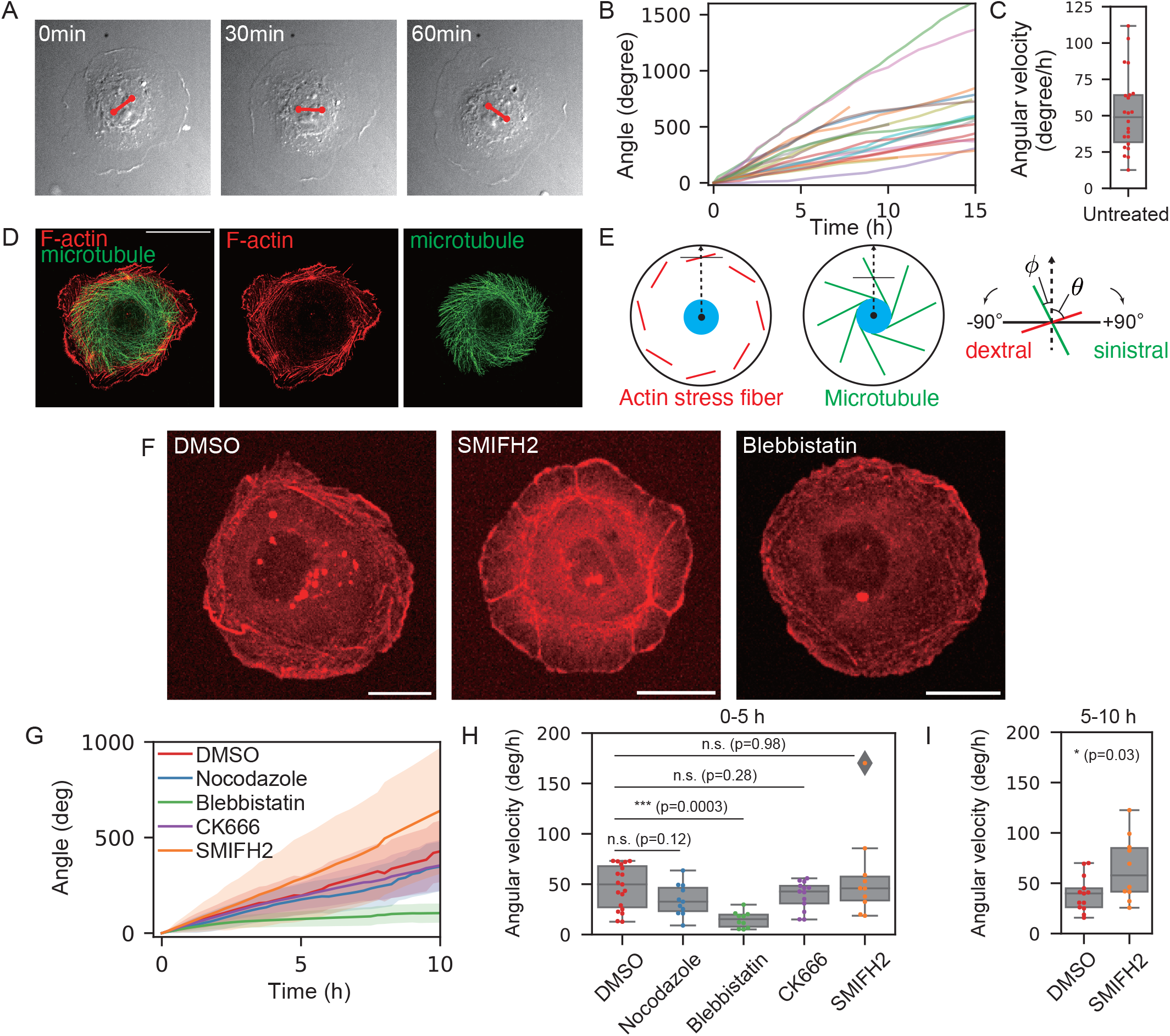
Chiral nuclear rotation and cytoplasmic flow in singly isolated Caco2 cells and the effect of inhibitors on it. (A) Rotating nucleus probed by the rotation of nuclear texture. The endpoints of the red line segments are the positions of tracked landmarks of the nucleus. (B) The cumulative angle of nuclear rotation plotted against time and (C) average angular velocity averaged over the first 10 hours (*n* = 22). Here, positive angle values indicate clockwise rotation. (D) Chiral cytoskeletal structure of F-actin (phalloidin) and microtubule (immunostaining). Scale bar: 20 μm. (E) Schematic diagram of the orientation of actin stress fiber and microtubule. (F) Snapshot images from the live image of actin dynamics in cells expressing Lifeact-RFP. Cells were treated with DMSO (0.2%), blebbistatin (1 μM), or SMIFH2 (40 μM). Scale bar: 20 μm. (G) The cumulative angle of nuclear rotation averaged over different cells plotted against time for different conditions: DMSO (0.2%, *n* = 19), nocodazole (50 μM, *n* = 11), blebbistatin (1 μM, *n* = 10), CK666 (200 μM, *n* = 13), SMIFH2 (40 μM, *n* = 10). The standard deviation is represented by shaded regions. (H) Angular velocity of nucleus under different conditions averaged over the first 5 hours of the time-evolution plot in (G). (I) Angular velocity of control and SMIFH2-treated cells averaged over the last 5 hours of the time-evolution plot in (I): DMSO (*n* = 13) and SMIFH2 (*n* = 10). p values were calculated using the Mann-Whitney-U test (* : *p* < 0.05, ** : *p* < 0.01, * * * : *p* < 0.001). Here, positive angle values indicate clockwise rotation.

The speed of nuclear rotation was about 50 degrees per hour on average (Fig. 1C). We measured the rotational speed by tracking unique points of the nuclear texture (Fig. 1A). The texture of the cytoplasm around the nucleus also showed a rotating motion, which indicates that the cytoplasm circulates in the same direction (Fig. 1A). Furthermore, we found that microbeads attached to the dorsal surface rotate (Movie S2), confirming that the dorsal membrane also rotates. The rotating motion of cells persists for more than eight hours until cell division occurs. After the cell division, cells form two-cell colonies, and then the nuclear rotation resumes. In this work, we focus on the rotating motion in singly isolated cells.

### F-actin and microtubules exhibit chiral patterns

We hypothesized that cytoskeletal molecules, such as F-actin and microtubules, are responsible for the circulating flow. To see the structure and dynamics of actin, we imaged live Caco2 cells expressing Lifeact-RFP (Movie S1). Actin bundles in the peripheral region of cells were tilted to form a dextral chiral pattern (Fig. 1D). Since each of these actin bundles associates with vinculin (Fig. S1), a focal adhesion protein, at their termini, we call them stress fibers [28]. We next studied how microtubules are organized in the cytoplasm during nuclear rotation. To this end, we visualized microtubules with the GFP-tagged microtubule-binding domain of ensconsin (EMTB-3XGFP [29]). Microtubules spread over the entire cytoplasmic region and exhibited a sinistral chiral pattern (Fig. 1D, Movie S3), consistent with the rotational direction of the nucleus. To summarize, actin bundles in the cell peripheral region and microtubules in the cytoplasm showed chiral patterns.

### F-actin and myosin-II are indispensable for chiral rotation

To investigate whether the cytoskeletons with chiral patterns drive the circulating flow, we performed live imaging of Caco2 cells expressing Lifeact-RFP with small-molecule inhibitors of cytoskeletal structures. When cells were treated with the actin polymerization inhibitor latrunculin A or F-actin stabilizer jasplakinolide, the shape of the cell periphery became rough and nuclear rotation stopped (Fig. S2, Movies S4 and S5, respectively), indicating that F-actin is necessary for the nuclear and cytoplasmic rotation. In contrast, disruption of microtubules by nocodazole did not affect the nuclear rotation (Figs. S2, S3, 1F and H, and Movie S6), which indicates that microtubules are not involved in the rotating motion.

To reveal which activities of actin are involved in the chiral rotating motion, we first investigated the role of Arp2/3-driven actin polymerization on the rotating motion, since a previous report has shown that it is involved in the chiral behavior of HFF [17]. When Caco2 cells were treated with the Arp2/3 complex inhibitor CK666 [30], lamellipodia at the cell periphery tended to shrink (Fig. S2, Movie S7), but the nuclear rotation was maintained (Figs. 1G and H), indicating that the Arp2/3 complex was dispensable for the rotating motion, in contrast to HFF where the Arp2/3 complex plays a role in cell chirality formation [17].

Next, we focused on the potential role of formin, a regulator of actin poymerization, as previous studies showed that this protein is involved in inducing the chirality of some cell types [7, 8, 17, 18, 31, 32]. To this end, we tested the effect of SMIFH2, a formin inhibitor [33], on the chiral pattern of Caco2 cells. When cells were treated with this inhibitor, chiral stress fibers mostly disappeared in their peripheral regions, but instead, another pattern of F-actin appeared (Fig. 1F and Movie S8). To investigate the distribution of F-actin more closely, we performed phalloidin staining and imaged cells in 3D (Fig. 2). Figure 2B shows that a subset of actin bundles became oriented in a radial direction, unlike the chiral pattern of stress fiber originally observed in the control cells (Fig. 2A). Furthermore, another population of F-actin was organized into a dense network or cluster with a concentric pattern (Fig. 2B). Intriguingly, in spite of these drastic changes in the spatial organization of F-actin, and the disappearance of the peripheral chiral stress fibers, the rotating motion was maintained (Figs. 1G, and H). Furthermore, as shown in Fig. 1I, we noticed that, while the rotating speeds of control and SMIFH2-treated cells were comparable in the first five hours of the observation window, the SMIFH2-treated cells rotated significantly faster than control cells, on average, in the second five hours: the rotating speed of control cells slightly decreased over time, while SMIFH2treated cells maintained or even slightly accelerated the rotating speed (Figs. 1H-I). These findings suggest that formins are not essential for the chiral rotation of nucleus in Caco2 cells, contrasted with previous observations [7, 8, 17, 18, 31, 32], and this inhibitor rather promoted the chiral rotation. Although we were not able to determine how SMIFH2 treatment induced F-actin reorganization, as this reagent is not strictly specific to formins [34], we used this inhibitor as a tool to investigate the mechanism of chiral rotational motion in further analysis.

**FIG. 2.**
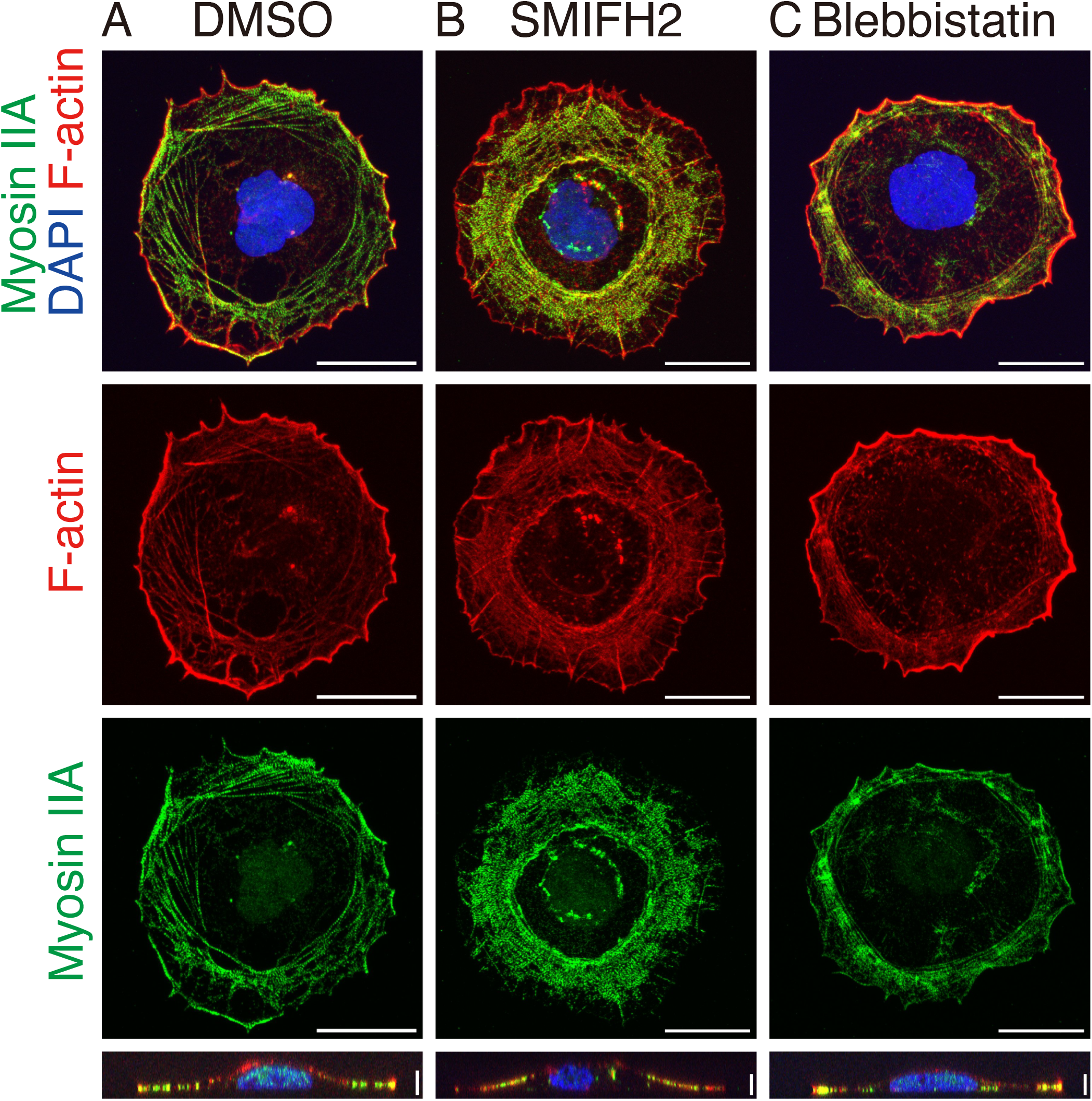
Organization of F-actin and Myosin IIA. (A) Control cells treated with DMSO show a chiral tilted pattern of F-actin and myosin II visualized by phalloidin and immunofluorescence with an antibody against myosin IIA, respectively. (B) SMIFH2 (40 *μ*M) treated cells show a concentric pattern of F-actin and myosin II. (C) The chiral tilted pattern of F-actin and myosin II is suppressed in cells treated with blebbistatin (1 *μ*M). The bottom panel shows vertical crosssections. Scale bars: 20*μ*m (horizontal) and 5*μ*m (vertical).

Previous studies have shown that the chirality in various cell types depends on the activity of myosin II, proposing that myosin II with F-actin generates chiral torque on a molecular scale [3, 35–37]. Therefore, we next investigated the role of myosin II in the chiral rotation by treating Caco2 cells with a myosin II inhibitor, blebbistatin. Under this condition, the chiral pattern of peripheral F-actin became less prominent (Figs. 1F, 2C and Movie S9), while the nuclear rotation was mostly suppressed (Figs. 1G and H). To confirm the role of myosin II, we depleted myosin IIA and/or myosin IIB heavy chains in Caco2 cells using siRNAs (Fig. S4B). Their depletion resulted in a significant reduction in the nuclear rotation (Fig. S4A). These results suggest that the activity of myosin II is required for the chiral rotational motion and also for the formation of a chiral pattern of stress fibers. On the other hand, our aforementioned results obtained using SMIFH2 indicated that the peripheral stress fibers forming a chiral pattern are not a critical component that drives the rotational motion, suggesting myosin II controls other subcellular mechanisms to induce the chiral rotational motion. In summary, both the activities of F-actin and myosin II are important for nuclear and cytoplasmic rotation.

### Super-resolution 3D imaging of actin and myosin-II

To investigate the roles of F-actin and myosin II in the chiral rotation further, we analyzed the distribution and dynamics of F-actin and myosin II in more detail, using control and SMIFH2-treated cells.

To this end, we performed a 3D super-resolution microscopy called expansion microscopy (ExM). We first observed the distribution of F-actin stained with phalloidin under control conditions (Fig. 3A). In the peripheral region of the cell, stress fibers showed a dextral swirling pattern (yellow in the left panel and bold lines in the right panel of Fig. 3A; dark red line Fig. 4I left) that are localized at the ventral side (yellow in Fig. 3A). Around the inner edge of the peripheral cytoplasmic zone having stress fibers, we detected another population of actin filaments, which are thinner than stress fibers, and do not exhibit obvious chiral orientation but are parallel to the tangential direction of the cell edge, which are distributed along the dorsal membranes of the cell (green in the left panel and dotted lines in the right panel of Fig. 3A; light blue line in Fig. 4I left). These observations suggest that the actin filaments along the dorsal membrane appear not to associate with other F-actin populations, unlike the previous observation on the cell chirality of HFF [17]. We also used antibodies to probe the distribution of myosin IIA (Fig. 3B). The distribution of myosin IIA is similar to that of F-actin, except for its striped pattern. Since we could not assess the colocalization of F-actin and myosin IIA filaments in the same cells in ExM due to a technical reason, we investigated their localization using conventional confocal microscopy (Fig. 2A and Movie S10). The confocal microscopy images indicate that F-actin and myosin IIA generally colocalize with one another in these specimens.

**FIG. 3.**
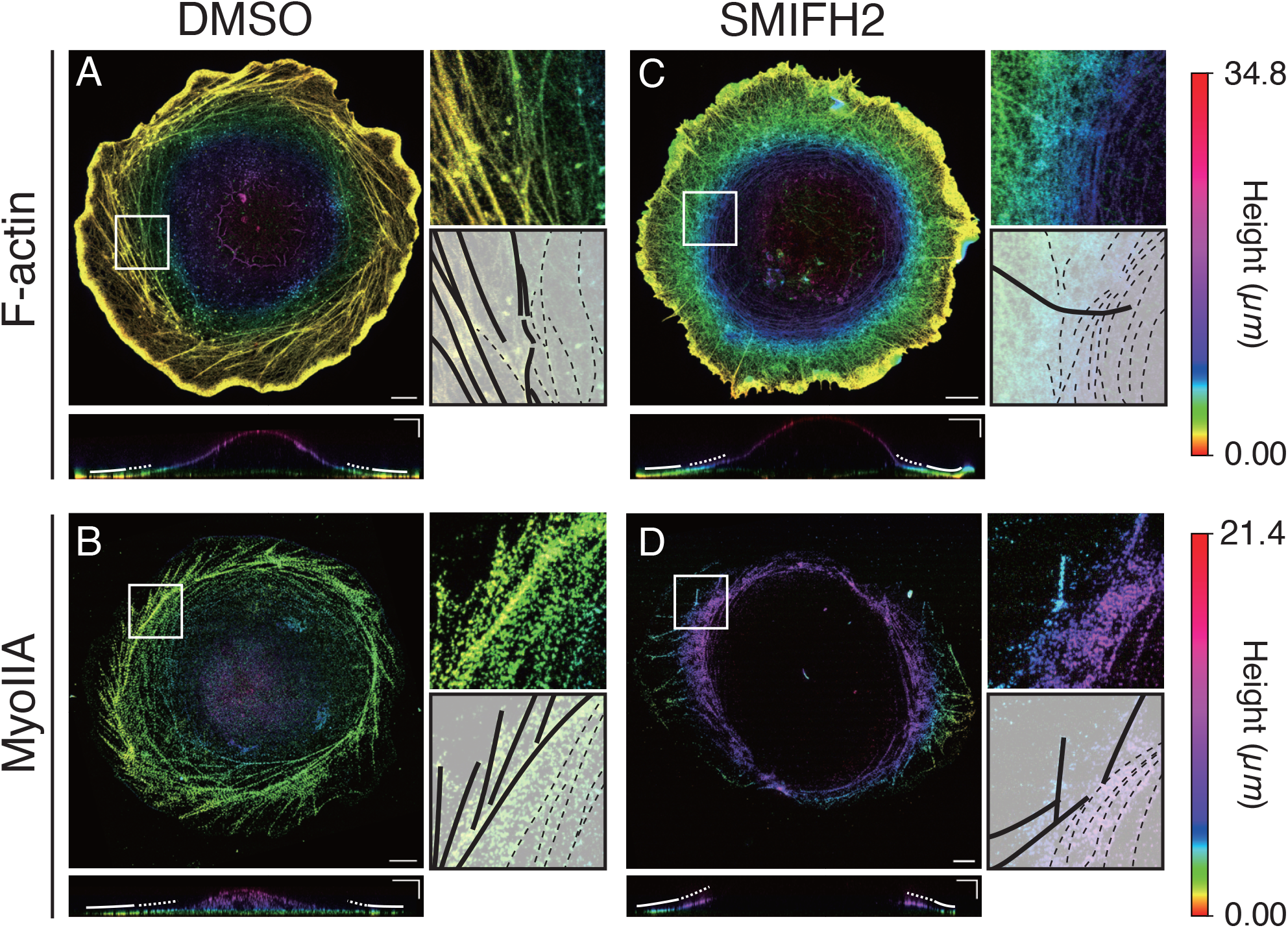
ExM imaging of F-actin and myosin IIA. Maximum intensity projection (MIP) images of F-actin (A, C) and myosin IIA (B, D) in DMSO (A, B) and SMIFH2 (C, D) treated cells. The color indicates the height along the *z*-axis, where the height was measured after the samples were swollen (color bar, right). Magnified views of the white boxes are shown in the right top panels, and corresponding outlines of F-actin are shown in the right bottom panels, where the bold and dotted lines indicate thick and thin fibers, respectively. The vertical cross-sections (*xz*) are shown in the bottom panels, where the bold and dotted lines indicate the peripheral and dorsal inner regions, respectively. Scale bars: 20 μm (horizontal) and 10 μm (vertical).

**FIG. 4.**
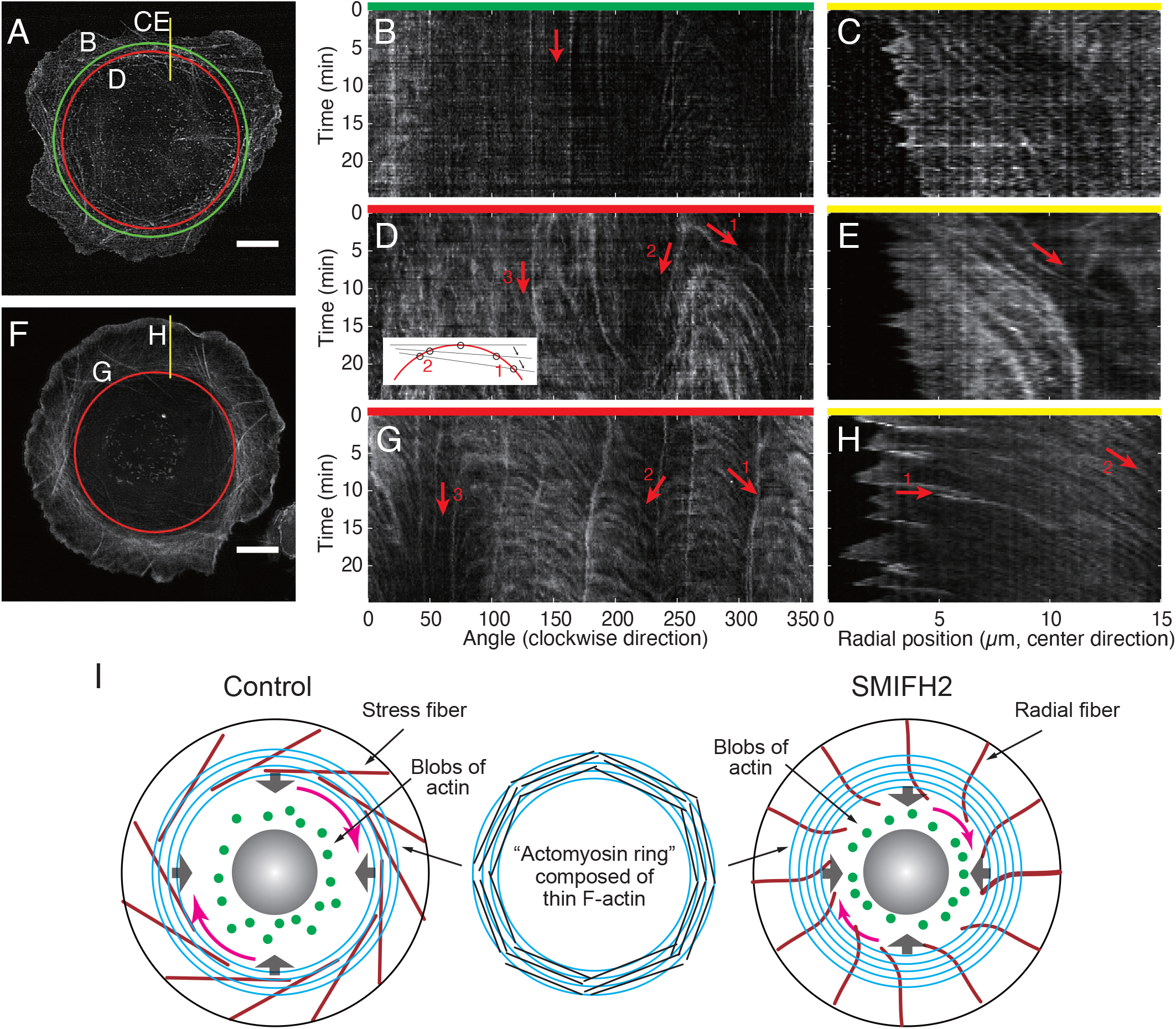
Dynamics of F-actin in Caco2 cells live-imaged by lattice light-sheet microscopy (LLSM). (A) Maximum intensity projection (MIP) image of Caco2 treated with DMSO. (B) Kymograph along the green circle in (A), obtained from a slice at *z* = 0. *z* = 0 is defined as the plane closest to the substrate. (C) Kymograph along the yellow line in (A), obtained from a slice at *z* = 0. (D) Kymograph along the red circle in (A), obtained from a slice at *z* = 0.5 *μ*m. inset: schematic diagram of F-actin (black lines) passing through the circle. (E) Kymograph along the yellow line in (A), obtained from a slice *z* = 0.5 *μ*m. (F) MIP image of Caco2 treated with SMIFH2 (40 *μ*M). (G) Kymograph along the red circle in (F), obtained from the MIP image. (H) Kymograph along the yellow line in (F), obtained from the MIP image. (I) Schematic diagram of F-actin structure in control and SMIFH2 treated cells. Thick stress fibers (dark red) were immobile, while the “actomyosin ring” (light blue), which consists of thin actin filaments, moved in centripetal and clockwise directions. Scale bar: 10 μm.

We next examined the distribution of F-actin and myosin II in the cells treated with SMIFH2 (Figs. 3C and D). As observed by conventional confocal microscopy, the chirally tilted actin stress fibers disappeared at the peripheral region, and instead thick F-actin bundles extended radially from the cell edge toward its center (bold line in Fig. 3C right; dark red line in Fig. 4I right). In the interior region, thin actin filaments were organized into a dense network with a concentric pattern, which were distributed at the dorsal side of cells (green in the left and dotted lines in the right panels in Fig. 3C; light blue line in Fig. 4I right). Myosin IIA exhibited a similar reorganization as seen in F-actin (Fig. 3D). As observed in control cells, confocal microscopy showed that F-actin and myosin IIA colocalize also in the SMIFH2-treated cells, particularly in the concentric actin clusters (Fig. 2B). Thus, ExM analysis revealed more detailed features of actomyosin distribution, particularly detecting its concentric orientation, located more inside the cell than the peripheral stress fibers. Additionally, we examined cells treated with blebbistatin by ExM, confirming the results obtained by the live imaging and the conventional immunostaining (Figs. 1F and 2C) that chiral stress fibers were greatly reduced after this treatment (Fig. S5).

### F-actin consisting of the “actomyosin ring” flows

To gain further insights into the role of the actomyosin system in the mechanism of intracellular rotation, we examined how F-actin behaves during the rotational process. To this end, we performed live imaging of Caco2 cells expressing Lifeact-mEmerald, using lattice lightsheet microscopy (LLSM). Since LLSM has a higher spatial resolution particularly in the z-direction compared to conventional confocal microscopy, we could identify the dynamics of F-actin in 3D more precisely. Figures 4B-E show kymographs of F-actin dynamics along different lines, drawn in Fig. 4A, at different heights *z* in control cells. In Figs. 4B and C, the kymographs along the green circle and the yellow line, which were analyzed at the ventral side, indicate that F-actin bundles in the peripheral region with the chiral tilted pattern are immobile on more ventral sides (arrow in Fig. 4B, see also Movie S11). Figures 4D and E show the kymographs along the red circle and the yellow line (drawn in Fig. 4A), respectively, at the height where the dorsal cell membrane exists. Rightward descending lines in the kymograph along the red circle indicate that the filaments move in a clockwise direction (Fig. 4D, arrow 1). There are also leftward descending lines that appear at the same time as the rightward descending lines appear but with different steepness (Fig. 4D, arrow 2). These pairs of lines indicate that the filaments are moving clockwise as well as centripetally (Fig. 4D inset). Furthermore, the kymograph along the yellow line (Fig. 4A) also indicates that the filaments are moving centripetally (Fig. 4E). To summarize, on the dorsal cell membrane, the concentric actin filaments move clockwise while also moving in centripetally (see also Movie S11): we hereafter call this concentric structure “the actomyosin ring” (Fig. 4I).

To see if the actomyosin ring is involved in driving the rotating flow, we estimated the spatial distribution of flow speed and orientation (velocity field) from the F-actin time-lapse images using particle image velocimetry (PIV) (Figs. 5, S6, and Movie S12). In the cytoplasmic region between the actomyosin ring and the nucleus, we did not detect clear actin filaments, but only found blobs of actin (Fig. 4A, and green dots in Fig. 4I). These blobs also circulate clockwise. From the velocity field inside the cells inferred by PIV, we calculated the angular component of the velocity with respect to the cell center and then converted it into the angular velocity, i.e. change in the angle per unit time. The spatial profile of the angular velocity (Figs. 5A and S6A-C) indicates that it is higher in the region where transverse fibers are present, rather than the region where the actin blobs are present, indicating that the driving force could be present in the region of the actomyosin ring.

**FIG. 5.**
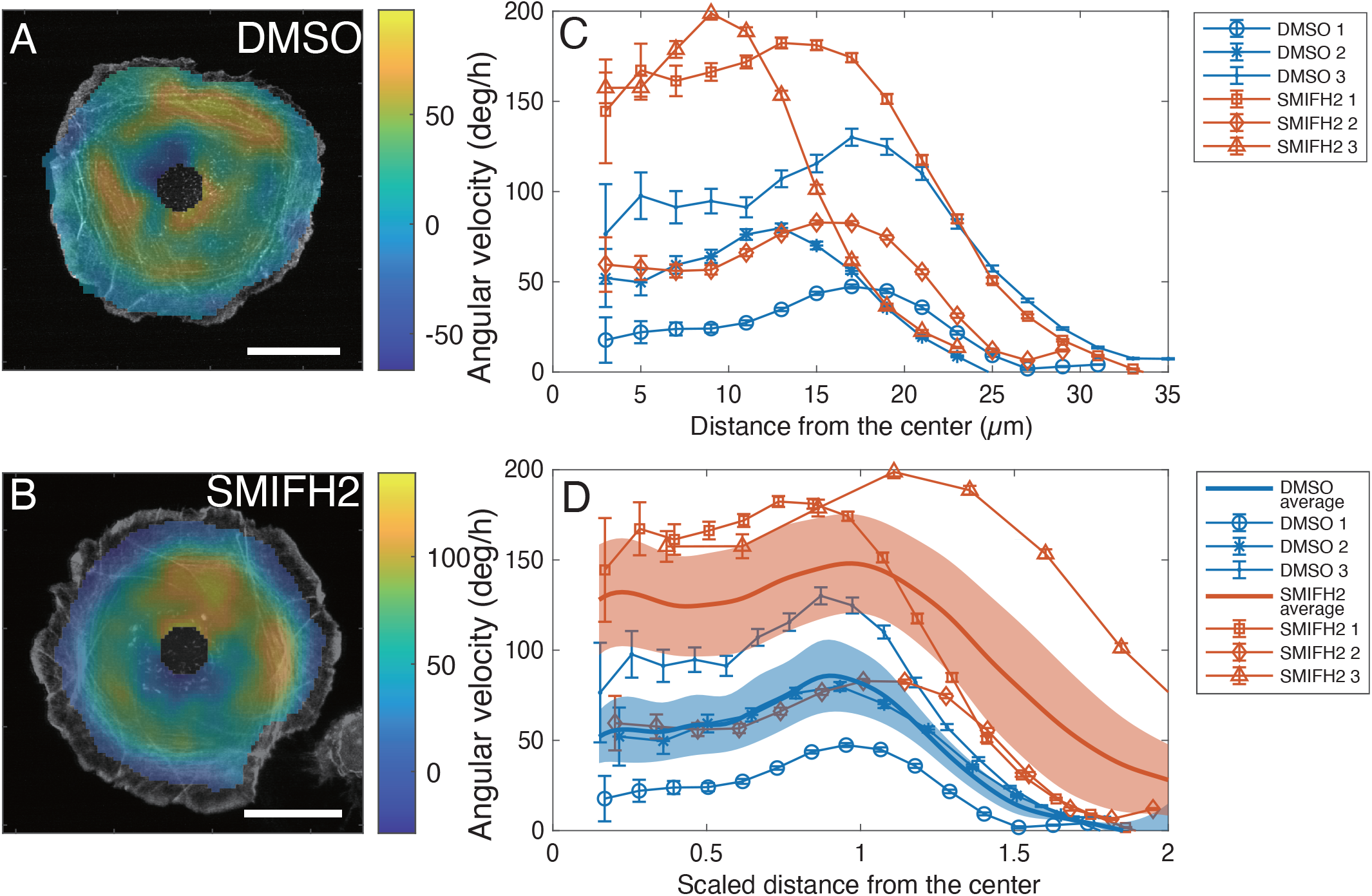
Angular velocity obtained by the PIV analysis. (A-B) Spatial profile of angular velocity (color code) obtained from the time-average of the PIV vector filed in a control cell (A: DMSO) or in a cell treated with SMIFH2 (B) superimposed on a snapshot F-actin image. Scale bar: 20 μm. (C) Average angular velocity as a function of the distance from the center. (D) Average angular velocity as a function of a distance scaled by the inner radius of the actomyosin ring of individual cells. Here, positive angular velocity indicates clockwise rotation. Sample averages for two conditions are indicated by the solid lines. Error bars and shaded areas represent standard errors of the means (SEM).

The angular velocity averaged over the angular direction was then plotted along the radial direction for control cells (Fig. 5C) and the peaks were found in the range from 10 to 20 μm. Since the size of the actomyosin ring varies from cell to cell, we manually determined the region of the actomyosin ring, and then replotted the angular velocity against the distance scaled by the inner radius of the actomyosin ring (Fig. 5D). We found that the peak positions are located around the scaled distance of one, which suggests that the driving force is present in the region around the actomyosin ring.

### Actomyosin ring flows in SMIFH2-treated cells

We also examined the dynamics of actin bundles in cells treated with SMIFH2, using live image data obtained by LLSM (Figs. 4F-H). In Figs. 4G and H, the kymographs along the red circle and yellow line (drawn in Fig. 4F), respectively, indicate that the actomyosin filaments organizing the ring move clockwise (Fig. 4G), and simultaneously flow centripetally (Fig. 4H), similar to the control condition. Additionally, we found that in contrast to the immobile stress fibers with a chiral pattern in control cells (dark red line in Fig. 4I left), F-actin bundles radially extending from the cell periphery in SMIFH2-treated cells tended to move passively in a clockwise direction at their proximal ends, although they seem to keep the anchorage of the distal ends to the cell edge (Fig. 4B, see also Movie S13 and dark red line in Fig. 4I right), implying that these radial F-actin bundles do not play active roles in the chiral motion of the actin ring. Thus, the concentric ring of flowing actomyosin filaments was detected also in the SMIFH2-treated cells, but showing modified features (Fig. 4I). Importantly, the ring developed more extensively after SMIFH2 treatment.

As in the control cells, the angular velocity inferred by PIV was high at the region around the inner edge of the actomyosin ring (Figs. 5B-D), which indicates that the driving force of the circulating flow is located in the region of the actomyosin ring as seen in the control cells.

We additionally observed another interesting phenomenon to support our idea. In a SMIFH2-treated cell that was imaged by a conventional fluorescence confocal microscope (LSM880, Zeiss), we, by chance, observed that fluorescent debris that seemed to attach to the actomyosin ring persistently circulated approximately three times as fast as the rotating speed of the nucleus: *∼* 400 degree/hour and *∼* 140 degree/hour for the debris and nucleus, respectively. (Movie S14, yellow and white lines, respectively). In the other cell observed by LLSM, we observed two fluorescent debris circulating in the area of the actomyosin ring, and in the cytoplasmic region between the actomyosin ring and the nucleus (Movie S15, yellow and red circles): the angular velocities of the circulating debris were *∼* 250 degree/hour and *∼* 190 degree/hour, respectively. Although we could not measure the rotating speed of the nucleus in the second cell because the nucleus was barely visible in the LLSM live image, the circulating speeds of the debris are more than two times faster than the typical nuclear angular velocity of SMIFH2-treated cells (Figs. 1G-I). These observations support the notion that the actomyosin ring generates a driving force for rotating the nucleus and cytoplasm. Note that, since the angular velocity estimated from the motion of debris was faster than that obtained from the PIV analysis, our PIV analysis for F-actin dynamics may underestimate the flow velocity.

In order to investigate the effects of SMIFH2 treatment on the rotational dynamics, we compared the angular velocity profiles inferred from the PIV between control and SMIFH2-treated cells. As shown in Fig. 5D, the angular velocity profiles averaged over samples indicate that the SMIFH2-treated cells tended to exhibit a faster speed compared to the control cells. This finding is consistent with the observation that SMIFH2-treated cells tended to exhibit a faster nuclear rotation, as shown in Fig. 1I. Additionally, the centripetal movement of the actin ring was also faster in the SMIFH2-treated cells than the control cells (Figs. S6H). These correlations between the enhanced development of the actin ring and the faster movement of cytoplasmic components strongly support the idea that the actomyosin ring generates a driving force for the chiral movement in Caco2 cells.

### A theoretical model of chiral cytoplasmic flow induced by the actomyosin ring

Our observations indicate the possibility that the actomyosin ring is a cell-scale structure that drives the chiral cytoplasmic flow. Since the actin filaments of the ring seemed not to have contact with the peripheral stress fibers in control cells, their rotational motion should be driven solely through its own mechanism, without relying on other structures, contrasted with the previous model that the contact between transverse fibers and radial fibers plays a role in establishing the cell chirality in HFF [17]. Then, how can a concentric pattern without an obvious cell-scale chiral structure generate chiral circulating flow? We here theoretically address this question.

We employ a theoretical framework of active chiral fluid [3, 35, 36], which has been proposed to describe the fluid dynamics driven by active chiral components. We model the actomyosin ring as an active chiral fluid driven by two active elements: 1) a force dipole originating from the contraction force of actomyosin and 2) a torque dipole generated when a bipolar myosin II filament rotates two antiparallel actin filaments to create a pair of counter-rotating vortex flows (Fig. 6A). By representing the orientation of actomyosin fibers as orientational field **p** and the fluid velocity as **v**, the hydrodynamic equation is described by the Stokes equation with the active contributions:

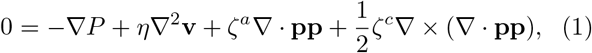

where *P* is the pressure satisfying the incompressibility condition ∇ · **v** = 0 and *η* is the fluid viscosity. Here, the terms with *ζ*^*a*^ and *ζ*^*c*^ are forces generated by the force dipoles (achiral) and torque dipoles (chiral), respectively. *ζ*^*a*^ and *ζ*^*c*^ represent the strength of the forces. The signs of *ζ*^*a*^ and *ζ*^*c*^ would be determined by the nature of force and torque generation at the molecular scale, independently of the cell-scale orientation of actomyosin. Considering that the actomyosin generates contractile force and right-handed torque as shown in Fig. 6A [25], the signs of the coefficients are *ζ*^*a*^ > 0 and *ζ*^*c*^ > 0. We hereafter assume *ζ*^*a*^ > 0 and *ζ*^*c*^ > 0 constant in space. We also assume the actomyosin filaments have a dipolar structure (Fig. 6A) so that Eq. 1 is invariant under **p** → −**p**. We, for convenience, represent the spatial variation of the density and order of the actomyosin by introducing an effective order parameter *S* as **p** = *S***n** (see Eq. 9), where **n** is a unit vector.

**FIG. 6.**
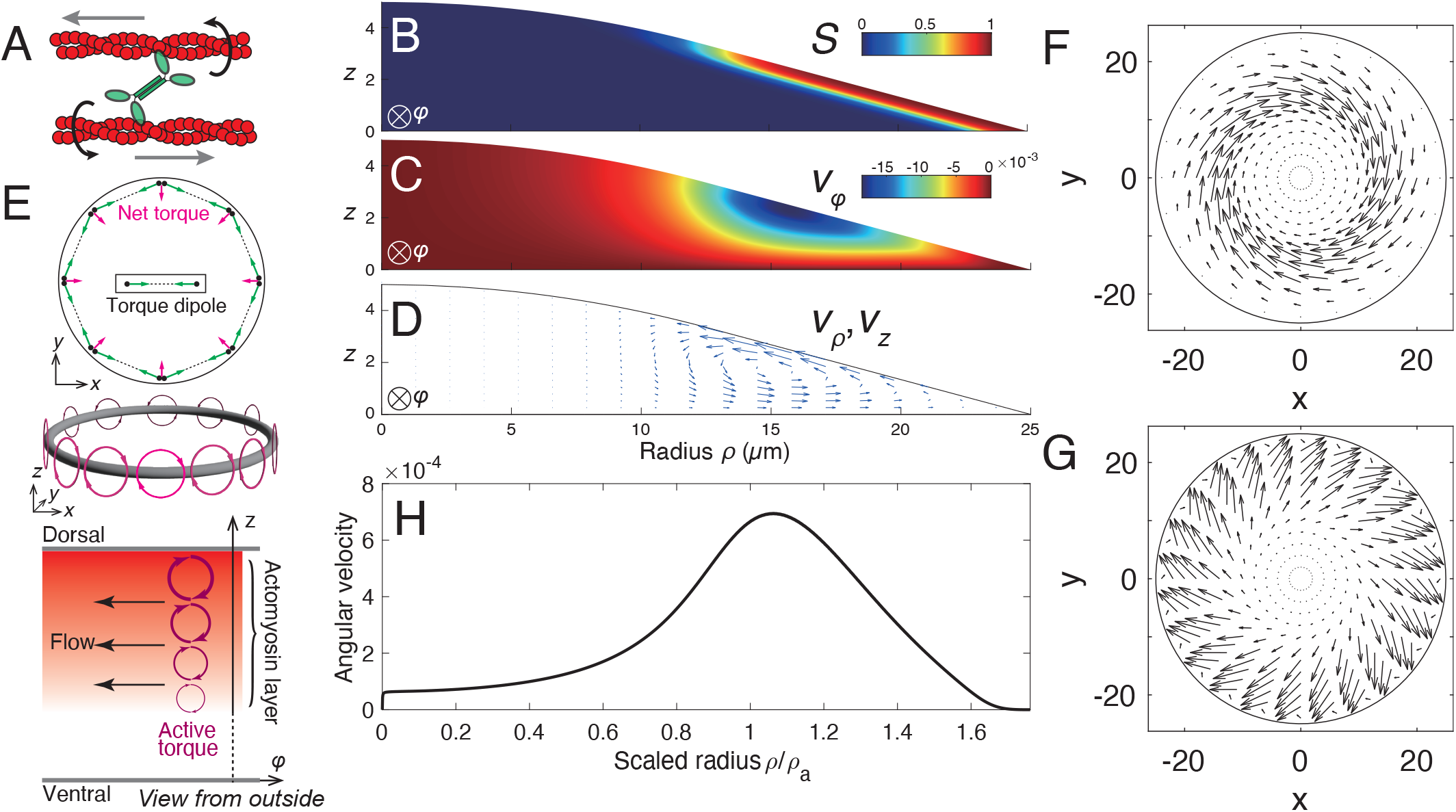
Theoretical model for the chiral cytoplasmic flow. (A) Actomyosin generates a force dipole and a torque dipole. (B-D) Numerical simulation of Eq. 1 assuming that the cell shape is axisymmetric around the cell center. (B) Actomyosin is distributed along the dorsal membrane with a concentric orientation. (C) Azimuthal velocity *v*_*φ*_ showing negative values indicating that the flow is generated in a clockwise direction. (D) Velocity in the radial *ρ*- and *z*-directions, (*v*_*ρ*_, *v*_*z*_), indicated by vectors. Circulating flow is generated in the *ρ*-*z* plane. (E) Top: A concentric orientational field on a ring generates net torque in the center direction (magenta arrow). Middle: Active torque (magenta clockwise arrow) generated by a concentric orientational field on a ring. Bottom: A gradient of active torque (magenta clockwise arrow) in the *z* direction, resulting in a rotational flow clockwise (black arrow). (F) Flow profile along the dorsal side showing an inward sinistral swirling pattern. (G) Flow profile at the ventral side showing an outward dextral swirling pattern. (H) Angular velocity averaged in the *z* direction plotted along the radial direction, showing a peak at around *ρ/ρ*_*a*_ = 1. Here, *ρ*_*a*_ is the leftmost position where *S ≥* 0.8.

We first numerically solved Eq. 1, assuming a cell with an axisymmetric geometry shown in Fig. S7. Based on experimental observation, we also assumed that actomyosin is distributed on the dorsal side, where the effective order parameter *S* is set to be positive reflecting the density distribution of actomyosin (Fig. 6B). Figure 6C shows the spatial profile of the azimuthal velocity *v*_*φ*_ in the vertical section of the cell. In the entire region, *v*_*φ*_ is negative, indicating that the flow is generated in a clockwise direction viewed from above. Thus, the numerical result shows that the concentric pattern of actomyosin can generate chiral cytoplasmic flow, and the direction is clockwise, consistent with our experimental observation. How can we understand the underlying mechanism behind the chiral cytoplasmic flow resulting from the concentric pattern of actomyosin? The active chiral term *ζ*^*c*^∇ × ∇ · **pp** in Eq. 1 can be interpreted as follows: the rotation of the axial vector field *ζ*^*c*^∇· **pp**, which is an active torque induced by chiral torque dipole, generates a force to induce a flow. For a concentric orientational field on a ring domain, the active torque *ζ*^*c*^∇· **pp** is generated as shown in Fig. 6E top. The actomyosin ring formed along the dorsal side in Caco2 cells is regarded as a stack of concentric orientational fields on a ring. In the region where actomyosin is present (*S* > 0), the concentration of actomyosin naturally increases with *z* which leads to an increase of the effective order parameter *S* with respect to *z* as indicated in Fig. 6B. For such an orientational field, the strength of the active torque increases as the height *z* increases (Fig. 6E bottom), forming a gradient of the active torque strength in the *z* direction. Consequently, the gradient generates a force in a clockwise direction as viewed from above (Fig. 6E bottom black arrow, see also the right-hand side of Eq. 10).

In the numerical simulation, we also investigated the spatial profile of *ρ* and *z* components (*v*_*ρ*_, *v*_*z*_) of the velocity field. Figure 6D shows that inward flow to the cell center occurs on the dorsal side, while the flow in the opposite direction occurs on the ventral side, resulting in the circulating flow in the *ρ*-*z* plane (Fig. 6D). This circulating flow is driven by the contractile force of actomyosin on the dorsal side. Due to the circulating flow in the *ρ*-*z* plane and the azimuthal flow, swirling flows appear on both dorsal and ventral sides (Figs. 6F and G, respectively). The sinistral swirling pattern at the dorsal side shown in Figs. 6F is driven by both centripetal and clockwise flows, which were indeed observed by LLSM shown in Fig.4. Interestingly, the dextral swirling pattern of the flow on the ventral side is consistent with the chiral pattern of the stress fibers on the ventral side (Fig. 1D). When the chiral flow was inhibited by blebbistatin, such a chiral pattern disappeared (Figs. 1F, 2C and S5), suggesting that the chiral tilted patterns are indeed formed in a flow-dependent manner. This implies that the dextral chiral pattern of the stress fibers is selforganized through alignment with the fluid flow on the ventral side.

Furthermore, we analyzed the radial distribution of the angular velocity and found that it exhibits a peak around the inner edge of the actomyosin ring *ρ ∼ ρ*_*a*_ (Fig. 6H), consistent with the PIV analysis of the experimental data (Figs. 5C and D). *ρ*_*a*_ is the inner radius of the actomyosin ring determined by the threshold *S ≥* 0.8. This result further supports the idea that the actomyosin ring drives the rotation of the nucleus.

### Depletion of dorsal actin and myosin stops the nuclear rotation

Our experimental observations and theoretical results suggest that the actomyosin ring located at the dorsal side of Caco2 cells plays a key role in the rotating motion. To further investigate this, we tested whether depletion of actomyosin at the dorsal side affected rotational motion. A previous study showed that the activation of RhoA by Rho Activator II (CN03), a specific RhoA activator, resulted in a decrease in apical stress fibers and an increase in basal stress fibers in vascular smooth muscle cells [38]. Based on this information, we aimed to decrease the ratio of the actomyosin at the dorsal to ventral sides in Caco2 cells using Rho Activator II. Remarkably, we observed a substantial increase in the thickness and number of actomyosin bundles at the ventral side, particularly beneath the nucleus, in Caco2 cells treated with Rho Activator II, while the dorsal actomyosin appeared to decrease significantly (Figs. 7A and C). The ratio of dorsal to ventral actomyosin was reduced significantly in the cell treated with Rho Activator II compared with control cells (Figs. 7D and E). We then performed live imaging of Caco2 cells treated with Rho activator II and found that the rotational motion of the nucleus ceased (Movie S16), supporting the idea that the dorsal actomyosin is crucial for driving the rotation. We also noticed that in cells treated with Rho activator II, the bundle of thick stress fibers was rearranged into a chordal pattern (Fig. 7C), and exhibited chiral motion (Movie S16), the mechanism of which remains to be understood. On the other hand, in cells treated with SMIFH2, the dorsal actomyosin appeared to increase, while the ventral actomyosin decreased (Figs. 7A and B). The ratio of dorsal to ventral actomyosin in cells treated with SMIFH2 tended to increase although the difference was not statistically significant (Figs. 7D and E). Taken together, our findings further support the idea that actomyosin at the dorsal side is crucial for driving rotation in Caco2 cells.

**FIG. 7.**
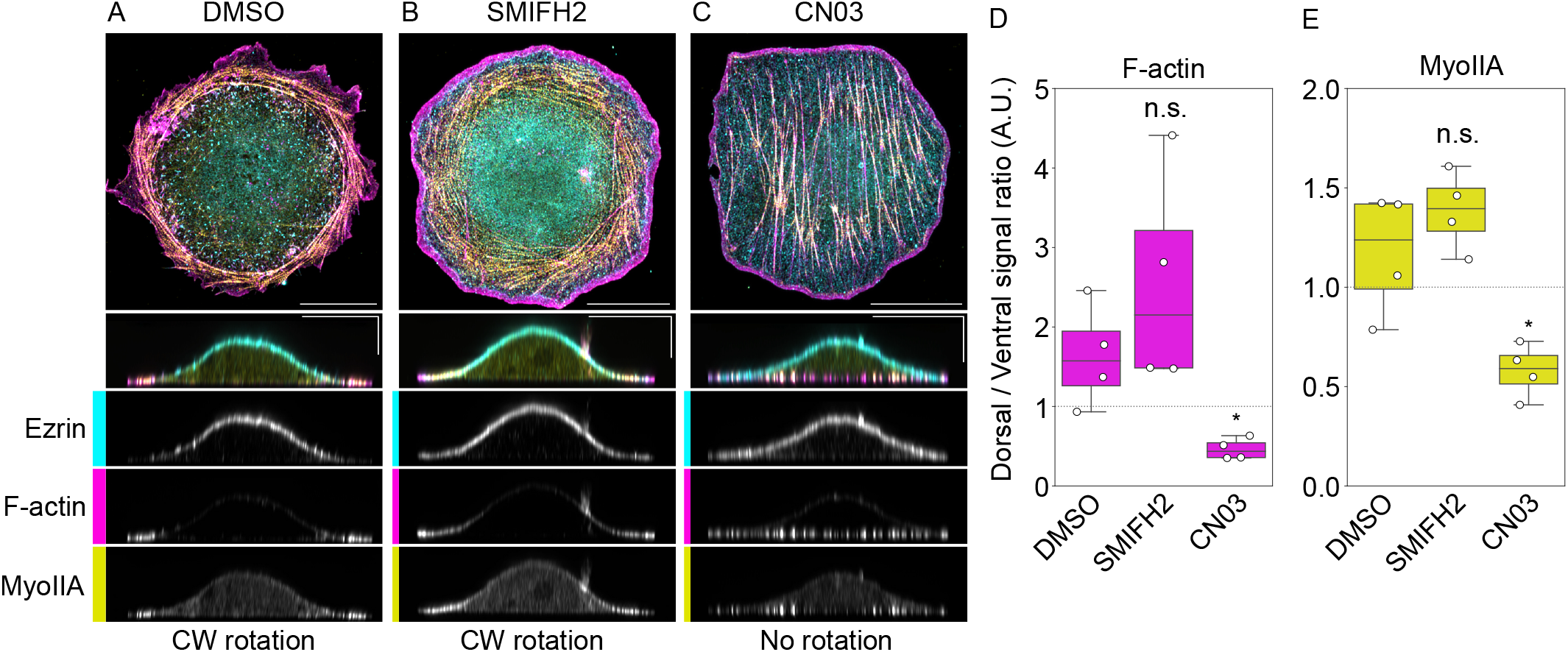
Depletion of dorsal actin and myosin by Rho activator II stops the rotating motion of nucleus. (A) Actin (magenta) and myosin II (yellow) showing localization with the dorsal marker Ezrin (cyan) in the DMSO treated cell. (B) SMIFH2-treated cell showing an increase in dorsal actomyosin and a decrease in ventral actomyosin. (C) Rho Activator II (CN03) treated cell showing a decrease in dorsal actomyosin and an increase in ventral actomyosin. Control (DMSO) and SMIFH2-treated cells showed clockwise (CW) rotation, while CN03-treated cells did show rotation. Scale bars: 20 μm (horizontal) and 5 μm (vertical). (D, E) Ratio of dorsal F-actin (D) and MyoII (E) to the ventral ones. p values were calculated using the Mann-Whitney-U test. * : *p* < 0.05, n.s.: *p* ≧ 0.05.

## DISCUSSION

In this study, we investigated the mechanism underlying cell-scale chiral dynamics, which is observed in Caco2 epithelial cells when cultured as a single cell. We found that Caco2 cells exhibited nuclear rotation and cytoplasmic circulation clockwise and these movements require actin and myosin II activities. High-resolution microscopy has revealed that the concentric actomyosin ring located on the dorsal side of the cells moves in a clockwise direction, leading us to hypothesize that this process may play a critical role in driving cytoplasmic flow. Previous studies using HFF proposed that radial actin fibers produce a force to drive the movement of the concentric actomyosin filaments (transverse fibers) through their connections [17]. In the case of Caco2 cells, however, we did not detect such radial fibers crossing the concentric actomyosin ring and our observations suggested that actomyosin ring by itself was moving in a clockwise direction through its own mechanism. To test this idea, we employed active chiral fluid theory [3, 35, 36], showing that the actomyosin localized under the dorsal membrane induces an active unidirectional fluid flow of the viscous cytoplasm and in turn nuclear rotation. Since the concentric pattern of actomyosin has no chirality at the cellular-scale, our theory indicates that the rotation of Caco2 cells is driven by the molecular-scale chiral mechanics of actomyosin rather than the cell-scale chiral orientation of actomyosin. It is also of note that we did not detect any visible cytoskeletal linkage between the nucleus and other cellular structures, another potential machinery for driving nuclear rotation. The nuclear rotation may be induced directly by the cytoplasmic circulating flow mediated by the friction between the nuclear surface and the cytoplasm. Microtubules and the peripheral stress fibers also exhibited chiral distribution patterns, but we obtained no evidence that they are involved in the chiral motion of the nucleus and cytoplasm. It is, therefore, possible that their chiral distribution was generated as a result of the above-mentioned mechanism.

Our experiments using an inhibitor and RNAi-mediated depletion have revealed that myosin II is involved in the chirality of Caco2 cells, which is consistent with previous studies showing the involvement of myosin II in the chiral behaviors of several types of cells [39]. Interestingly, while some of these studies concluded that formins are essential for breaking the chiral symmetry [7, 8, 17, 18, 31, 32], our results showed that the rotational speed of Caco2 did not decrease but even slightly increased when treated with SMIFH2, a known inhibitor of formins, suggesting that formins are not required for Caco2 cell chirality. Curiously, a previous study showed that SMIFH2 inhibits the centripetal movement of myosin II filaments in HFF [34], simultaneously demonstrating that this inhibitors also inhibits myosins, which apparently contradicts our observation. However, the same group also reported that SMIFH2 facilitated the centripetal movement of actin and myosin filaments when rat embryo fibroblasts were used [40], similar to our present observations. These reports suggest that how cells respond to SMIFH2 depends on their types. In the case of Caco2 cells, it is likely that the observed effects of SMIFH2 on actomyosin dynamics were not attributed to its potential inhibition of myosin II, although how this inhibitor induced the observed reorganization of actin and myosin filaments in Caco2 cells remains unknown, as SMIFH2 seems to have multiple targes [40], t.

In Caco2 cells treated with SMIFH2, the actomyosin ring became more visible than in the control cells (Figs. 2, 3 and 7). Cells treated with SMIFH2 tended to show a trend of actin and myosin shifting from the ventral side to the dorsal side compared to control cells, although not statistically significant (Fig. 7DE). This might be due to a decrease in the formation of stress fibers on the ventral side, followed by a shift of free actin and myosin II to the dorsal side, which may lead to an increase in the formation of actomyosin ring at the dorsal side. In contrast, treatment by Rho Activator II increased the stress fiber formation at the ventral side, which can induce a shift of actin and myosin II to the ventral side, leading to a decrease in the formation of actomyosin ring at the dorsal side (Fig. 7DE). These considerations provide support for the model that dorsal actomyosin constitutes the driving force behind the rotational motion, as the rotation tended to increase or decrease in SMIFH2 or Rho Activator II-treated cells, respectively.

In a previous study [39], an achiral active fluid model was proposed to explain the rotation of the nucleus driven by actomyosin. In the theory, the concentric orientational order of actomyosin becomes unstable due to the spontaneous chiral symmetry breaking induced by the contractility of actomyosin, and then a chiral orientational order emerges to drive a unidirectional fluid flow to rotate the nucleus. Since there is no intrinsic chirality in the model, either clockwise or counter-clockwise rotation is selected with equal probability. In contrast, in our theoretical model, we considered the intrinsic chirality of the actomyosin gel in order to explain our experiments where the rotational direction is always clockwise. Although our model is consistent with our experimental observations, there is still a limitation. We assumed a concentric orientational order of actomyosin. However, it remains to be elucidated how the concentric order is formed [41, 42], and how stable the structure is when the actomyosin has intrinsic chirality.

Several types of cells have been reported to exhibit chiral nuclear rotation. Zebrafish melanophores exhibit a counterclockwise nuclear rotation “from basement view” [16], which is the same as our observation where the nucleus rotates in a clockwise direction viewed from the dorsal side. The study reported that actin plays a pivotal role in the chiral rotational motion. In the case of singly isolated MDCK cells embedded in a 3D culture, nuclei as well as whole cells exhibited rotations in either direction with a bias to the counterclockwise direction [19]. Furthermore, a weak inhibition of actin polymerization reversed the bias in the rotational motion, and an actin-binding protein *α*-actinin-1 regulates the direction of chiral rotation [19], similar to the case of HFF [17, 18]. In the case of C2C12 myoblasts, whether actin filaments were organised or disorganised was shown to correlate with chirally biased nuclear rotation [43]. In contrast to HFF [17, 18], the direction of nuclear rotation of Caco2 is opposite and the chirality of Caco2 does not require the activity of Arp2/3 complex (Figs. 1 and S2) nor physical interactions between the actin ring and other actin structures such as radial fibers. Thus, the mechanism of cell chilarity formation seems to be different between cell types, such as HFF and Caco2 cells. Whether there is a common principle behind the chiral nuclear rotation or whether they are caused by different mechanisms remains to be clarified in future studies.

It has been suggested that the left-right asymmetry at the tissue level originates from chirality at the cellular level. However, it remains unclear how cell chirality coordinates to induce left-right asymmetry in multicellular organisms. Our previous theoretical studies showed that a tissue-scale asymmetry, such as a spatial gradient in the strength of cell-scale torque generation, is necessary for the tissue-scale left-right asymmetry to arise from cell chirality [12, 21, 22]. Hence, it will be necessary to investigate how cell chirality and tissue-level asymmetry coordinate to understand left-right asymmetry in organs and the body. Caco2 is an epithelial line, which can form multicellular layers. We thus hope that investigating the coordination between cell chirality and tissue-level chirality using cells, such as Caco2, which exhibit a clear individual chirality, is a promising approach to reveal the principles of chiral morphogenesis.

## ACKNOWLEDGMENTS

This work was supported by Kakenhi grant 19K16096 (TY) 23K14186 (TI), 22H05170, 23H02455 (TS), and JST CREST Grant JPMJCR1852 (TS) and the core funding at RIKEN Center for Biosystems Dynamics Research. TY and TI were suported by Grant-in-Aid for JSPS Fellows 18J01239 (TY) and 22KJ3145 (TI). We thank H. Saito, T. Kato, and H. Hamada for the advice and support during experiments, M. Hayakawa, G. Ogita, B. Bhattacherjee, R. Nishizawa, K. Kawaguchi, K. Adachi, Y. Fukai, R. Cerbus, T. Kato, S. Fürthauer, Y. H. Tee, and A. D. Bershadsky for fruitful discussions, and W. Kimura for sharing reagents.

## AUTHOR CONTRIBUTIONS

Conceptualization, T.Y., T.I., M.Takeichi, and T.S.; Investigation, T.Y., T.I., Y.M-K., S.H., N.T., M. Tarama, M. Takeichi and T.S.; Methdology, T.Y., T.I. and T.S.; Formal analysis, T.Y., T.I. and T.S.; Funding acquisition, T.Y., T.I., and T.S.; Writing – original draft, T.Y., T.I. and T.S.; Writing – review & editing, T.Y., T.I., Y.M-K., M. Tarama, M. Takeichi and T.S. Supervision, T.S.

## DATA AND CODE AVAILABILITY

Source data for all graphs and original code for the numerical integration of the model have been deposited and is publicly available at https://doi.org/10.5281/zenodo.8254364.

## MATERIALS AND METHODS

### Cell cultures and transfection

Caco2 (ATCC) cells were cultured in DMEM/Ham’s F-12 (FUJIFILM Wako Pure Chemical Corporation, 048-29785) supplemented with 10% fetal bovine serum (SIGMA, F7524, Lot. BCBV 4600) and 1% penicillin/streptomycin (nacalai, 26253-84) at 37°C, 5% CO2 on collagen type I coated Dish (60 mm, IWAKI, 4010-010). For live-imaging of actin, we used LifeactRFP-transfected Caco2 cells which were established from Caco2 (ATCC) in [27]. To live-image the microtubule dynamics, we transiently transfected Caco2 cells with EMTB-3XGFP using Lipofectamine LTX Reagent with PLUS Reagent (Invitrogen, 15338100), according to the manufacturer’s protocol. EMTB-3XGFP was a gift from William Bement [29] (Addgene plasmid # 26741 ; http://n2t.net/addgene:26741; RRID:Addgene 26741) For Lifeact-mEmerald (pLVSIN-EF1a-Lifeact-mEmerald-IRES-pur), mEmerald-Lifeact-7 was inserted into pLVSIN-EF1a-IRES-pur at the BamHI and NotI sites by In-Fusion. The construction of pLVSIN-EF1a-IRES-pur has been described previously [44]. mEmerald-Lifeact-7 was a gift from Michael Davidson (Addgene plasmid # 54148 ; http://n2t.net/addgene:54148 ; RRID:Addgene 54148) For protein depletion, cells were transfected with Stealth siRNA using Lipofectamine RNAi MAX (Invitrogen). The following siRNAs were used: MYH9HSS106871 for myosin IIA; MYH10HSS106875 for myosin IIB; Negative Control Med GC Duplex (#10002823) for negative control, and we examined the effects of RNA interference at five days.

### Immunoblotting

Cells were collected in RIPA buffer containing 1x cOmplete EDTA-free protease inhibitor (Roche, 05056489001) and were then mechanically lysed by passage through a 27G needle on ice. Samples were boiled with 10% 2-mercaptoethanol for 3 minutes and separated by SuperSep 7.5% SDS-polyacrylamide gel electrophoresis (Fujifilm, 198-14941) at 250 V / 30 mA for 70 minutes, and subsequently transferred to a 0.45 *μ*m pore size PVDF membrane (Cytiva, 1080682) at 250 V / 300 mA for 90 minutes in an ice bath. The PVDF membrane was blocked with Blocking One (Nacalai Tesque, 03953-95) at 4°C overnight, and incubated with appropriate primary antibodies for 60 minutes at room temperature (RT). After washing the membrane with Tris-buffered saline with 0.1% Tween 20 (TBST), it was incubated with appropriate HRP-conjugated secondary antibodies for 60 minutes at RT. The membrane was then washed with TBST, followed by enhanced chemiluminescence (ECL) detection (Bio-Rad, 1705060) using LAS-3000 mini (Fujifilm) for image acquisition. Signal intensity was analyzed using Gel Analysis tool in ImageJ Fiji [45]. The following antibodies were used: rabbit anti-Myosin IIA (Sigma-Aldrich, M8064, dilution 1:1000); rabbit anti-Myosin IIB (Cell Signaling Technology, 8824, dilution 1:1000); mouse anti-GAPDH (Santa Cruz Biotechnology, sc-166574, dilution 1:1000); HRP-conjugated goat anti-rabbit IgG (Invitrogen, T20926, dilution 1:1000); HRP-conjugated goat anti-mouse IgG (Invitrogen, T20912, dilution 1:1000). As a protein ladder marker, Precision Plus Protein Dual Color Standard (Bio-Rad, 1610374) was used.

### Live cell imaging

For live-image data in Figs. 1, S2 and S3, we used an inverted fluorescence microscope (Olympus, IX-81) equipped with a spinning disk confocal imaging unit (Yokogawa, CSU-X1), a 60x/1.35 oil immersion objective (Olympus, UPLSAPO60XO), and a 561 nm laser (Coherent, Sapphire LP) for RFP excitation or a 488 nm laser (Coherent, Sapphire LP) for GFP excitation. We used a 40×/1.35 oil-immersion objective (Olympus, UApo/340) for the microtubule snapshots in Fig. S3. The cells were seeded sparsely on a collagen type I coated glass-based dish (IWAKI, 4970-011), and incubated in a stage-top incubator (Tokai Hit) at 37°C with 5% CO2 during liveimaging.

We took fluorescence images with multiple z-stacks (number of slices: 7 and Δ*z*: 0.5 *μ*m) by EMCCD (Andor Technology, iXon+) every 15 min, and then made maximum intensity Z projections. For the microtubule snapshots in Fig. S3, we applied a different condition (number of slices: 5 and Δ*z*: 1 *μ*m).

For the inhibitor experiments, the following inhibitors were used: latranculin A (SIGMA, L5163-100UG); Jasplakinolide (Toronto Research Chemicals Inc, J210700); Nocodazole (SIGMA-Aldrich, M1404); CK666 (SIGMA-Aldrich, SML0006-5MG); SMIFH2 (Wako, 4401/10); Blebbistatin (SIGMA, B0560-1MG); Rho Activator II (Cytoskeleton, Inc., Cat. #CN03). We started live-imaging about 2-3 hours after seeding cells, and added the inhibitors about 40 min before the live-imaging.

When we observed the dynamics of the beads attached to the dorsal membrane of the cells, we used 2 *μ*m carboxylate-modified beads (Invitrogen, F8887), and live-imaged the dynamics using the DIC channel of the microscope (Olympus, IX-81).

### Analysis of cell rotation

We quantified the rotational behaviors of cells by manually tracking the dynamics of two nucleoli of each cell on DIC images (Segmentation Editor, Fiji). We defined the rotational angle of the cell by that of the line connecting the two nucleoli, and analyzed the rotational dynamics using Python. In Figs. 1A and B, we defined the initial time point of the measurement of the nuclear rotation by the time when we started the live-imaging. In the inhibitor experiments in Fig. 1, we determined the initial time point of the measurement as the time 5 hours after the initiation of live-imaging, accounting for the time lag needed for the inhibitors to exert their effects.

### Immunofluorescence antibody staining and microscopy

Cells were seeded on collagen type I coated cover slips (Neuvitro Corporation, NEU-H-12-COLLAGEN-45) and treated with the inhibitors. After 8 hours, the cells were fixed with 2% PFA in PBS(-) for 10 min, permeabilized with 0.25% Triton X-100 in PBS(-) for 10 min, blocked with 3% BSA in PBS(-) for 30min. Then, we incubated cells with primary antibodies (2 hours), secondary antibodies (1 hour) and phalloidin (30 min) in a blocking buffer (3% BSA in PBS(-)). After washing with PBS(-) three times, the samples were mounted with a mounting medium with DAPI (Vector Laboratories, VECTASHIELD, H-1200). All the processes were performed at room temperature.

We used rabbit anti-Myosin IIA (Sigma-Aldrich, M8064, 1:1000 for IF), mouse anti-Vinculin (Sigma, V9131, 1:200 for IF), and mouse anti-Ezrin (Abcam, ab4069, 1:1000 for IF) as the primary antibodies and Alexa Fluor 488 goat anti-rabbit IgG (Sigma-Aldrich, 11034, 1:1000 for IF), Alexa Fluor 488 goat anti-mouse IgG (Invitrogen, A11029, 1:1000 for IF), and Alexa Fluor 647 donkey anti-mouse IgG (Sigma-Aldrich, AP192SA6, 1:1000 for IF) as the secondary antibodies, respectively. For actin staining, we used Alexa Fluor 568 phalloidin (Invitrogen, A12380, 1:400).

To analyze the sample, we took fluorescence images with multiple z-stacks (Δ*z*: 0.32 *μ*m) using a laser scanning confocal microscope (Zeiss, LSM880) equipped with Plan-Apochromat 63x/1.4 Oil DIC M27. Images were processed with Fiji.

### Expansion Microscopy

Protein-retention Expansion Microscopy (ExM) was carried out as described previously [46]. Cells were cultured on collagen type I (SIGMA, C8919-20ML) coated cover slips and treated with the inhibitors. Fixation, permeabilization and blocking were performed as described above. To visualize F-actin, the cells were stained with Alexa Fluor 488 phalloidin (Invitrogen, A12379, 1:400) and next stained for 60 min with a rabbit anti-Alexa Fluor 488 antibody (abcam, ab150077, 1:500) as the primary antibody and Alexa Fluor 488-conjugated goat anti-rabbit (1:1000) as the secondary antibody [47]. After washing, cells were incubated with 100 *μ*g/mL of 6-((Acryloyl)amino)hexanoic acid, succinimidyl ester overnight at room temperature in the dark. Samples were incubated in gelation solution (8.6% (w/w) sodium acrylate, 2.5% (w/w) acrylamide, 0.15% (w/w) N,N’-methylenebisacrylamide, 2 M NaCl, 1 x PBS, 0.1% TEMED, 0.1% ammonium persulfate) for 5 min on ice. Gelation was allowed to proceed at room temperature for 1 hour. The gel and a cover slip were removed with tweezers and incubated with digestion buffer (0.5% Triton X-100, 1 x TE buffer, 1 M NaCl, 8 unit/mL Proteinase K) overnight at room temperature in the dark. The gels were removed from the digestion buffer and placed in 50 mL of Milli-Q water. Water was exchanged three times every 30 minutes. Most of the gels expanded to about 4.5 times their original size. Gels were placed on a poly-L-lysine (Sigma-Aldrich, P4707) coated glass bottom dish, and fluorescence images were taken by a laser scanning confocal microscope (Zeiss, LSM880).

### Lattice Light Sheet Microscopy (LLSM) and image processing

The LLSM was home-built in the Kiyosue laboratory at RIKEN Center for Biosystems Dynamics Research following the design of the Betzig laboratory [48] under a research license agreement from Howard Hughes Medical Institute. Electric wiring was performed at RIKEN Advanced Manufacturing Support Team. Metal Parts were processed by Maeda Precision Manufacturing Ltd. and Zera Development Co. To create a lattice light sheet, a dithered square lattice was used through a spatial light modulator (Fourth Dimension Displays) in combination with an annular mask with 0.55 out and 0.44 inner numerical apertures (Photo-Sciences) and a custom NA 0.65 excitation objective (Special Optics). Images were acquired through a CFI Apo LWD 25XW 1.1-NA detection objective (Nikon) and a scientific sCMOS camera, Orca Flash 4.0 v3 (Hamamatsu Photonics). Caco2 cells expressing Lifeact-mEmerald were seeded on a collagen-coated coverslip 3h before imaging. During imaging, cells were maintained in DMEM/Ham’s F-12 (FUJIFILM Wako Pure Chemical Corporation, 048-29785) supplemented with 10% fetal bovine serum (SIGMA, F7524, Lot. BCBV 4600) at 37°C, 5% CO2. For live imaging of Lifeact-mEmerald, a 488-nm laser (MPB Communications) and a long-pass emission filter BLP01-488R-25 (Semrock) were used. Image stacks were collected with a 200 nm step size between planes with 10 msec per plane exposure time and 14.8 sec time interval. After the acquisition, images were deskewed and deconvolved using LLSpy. After deskew processing, the voxel pitch was 0.104×0.104×0.103*μ*m.

### Particle image velocimetry (PIV) analysis

PIV analysis was performed using PIVlab [49] for the time-lapse images obtained by LLSM. The velocity vector fields were calculated using a multi-grid interrogation (64 × 64, 32 × 32, and 16 × 16 pixel sizes of interrogation windows with 50% overlap each). PIV was performed in the masked area, and the masked area was determined by thresholding the time-integrated image. Using the velocity vector field (*v*_*x*_, *v*_*y*_), we first calculated azimuthal (*v*_*φ*_) and radial (*v*_*ρ*_) velocities as 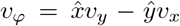 and 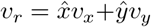, where 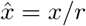 and ŷ = *y/r* with (*x, y*) being the position from the center of cell and 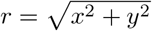. Here, the cell center was taken as the *xy*-coordinate of the highest position of the unimodal-shaped cell. From the DIC image, the nucleus always rotates around the center of the nucleus, with the highest position of the cell just above the nucleus center. Then, the angular velocity *ω* was obtained as *ω* = *v*_*φ*_*/r*. The temporal averages of *v*_*φ*_ and *ω* are shown in Fig. S6 for all samples analyzed (control cells (DMSO)(Figs. S6A-C) and cells treated with SMIFH2 (Figs. S6D-F)). The angular averages of *v*_*φ*_ (Fig. S6G), *v*_*r*_ (Fig. S6H) and *ω* (Fig. 5C) were obtained from their temporal averages at each spatial point. These averages were plotted against the radius scaled by the inner radius of the actomyosin ring in Fig. 5D. We here manually identified the inner edge of the actomyosin ring and calculated the inner radius for each cell.

### Theoretical model

We here describe a brief derivation of our 3D model [35]. We assume a low Reynolds number limit, a steady state, and incompressibility. In the theory of active chiral fluid, the momentum conservation is represented as:

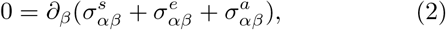

where 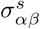 and 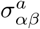 are the symmetric and asymmetric parts of the deviatoric stress, respectively. 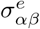 is Ericksen stress (hydrostatic stress). The indices *α, β*, and *γ* denote the three Cartesian coordinates *x, y*, and *z*. The constitutive equations of the deviatoric stress are given:

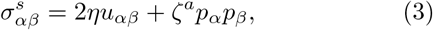

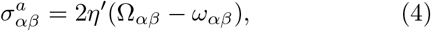

where *u*_*αβ*_ = (*∂*_*α*_*v*_*β*_ +*∂*_*β*_*v*_*α*_)*/*2 and *ω*_*αβ*_ = (*∂*_*α*_*v*_*β*_ −*∂*_*β*_*v*_*α*_)*/*2 is the strain rate and the vorticity. Ω_*αβ*_ is the spin rotation rate describing the intrinsic rotation rate of local volume elements. *η* and *η*^*′*^ are viscosity coefficients, and *ζ*^*a*^ is a coefficient of the achiral active stress. We here only consider anisotropic contributions of active terms allowed in a chiral nematic active fluid for simplicity. Also, in this study, since we assume that the orientational field **p** is fixed to be a concentric pattern, we omit the terms that derive from the molecular field. We thus define 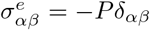, where *P* is the pressure serving as a Lagrange multiplier to satisfy the incompressibility.

The angular momentum conservation is given by the following equation:

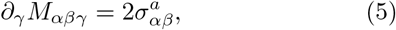

where *M*_*αβγ*_ is the angular momentum flux. The constitutive equation is written as:

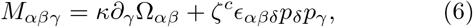

where *κ* is a dissipative coefficient and *ζ*^*c*^ is a coefficient of the active chiral stress which reflects the symmetry of the torque dipole represented in Fig. 6, which is called nematic chiral rod motor [35]. *ϵ*_*αβγ*_ is the Levi-Civita symbol.

We derive the following equation of motion from Eqs. 2-6,

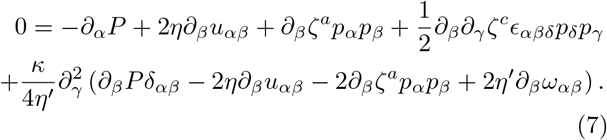

In the final term of Eq. 7, the length scale 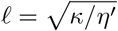 is a characteristic molecular scale. Since we consider the hydrodynamics at the cell scale, we take the limit of *ℓ* → 0 and omit the final term. Finally, applying the incompressibility condition *∂*_*γ*_*v*_*γ*_ = 0, we obtain the final form:

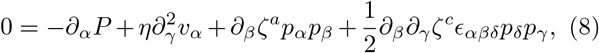

which is equivalent to Eq. 1.

In the numerical simulations, for simplicity, we suppose that the cell is axisymmetric as shown in Figs. 6 and S7. Based on the experimental observations, we consider that the actomyosin bundles align along the circumferential direction: the concentric pattern of the actomyosin ring. We here represent the orientational order **p** of the actomyosin in the cylindrical coordinate (*ρ, φ, z*). Since **p** is aligned in the circumferential direction, **p**(*ρ, z*) is given in the cylindrical coordinate by

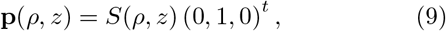

where *S*(*ρ, z*) is the effective strength of the orientation of the actomyosin and takes a finite value in the domain where the actomyosin ring is present. Since we did not see any specific orientational order in the direction of the cell height at least at our imaging resolution, we considered the orientation of the actomyosin bundle to be parallel to the substrate, and the *z* component of **p**(*ρ, z*) is zero. In the cylindrical coordinate, Eq. 1 of motion for the fluid velocity **v** = (*v*_*ρ*_, *v*_*φ*_, *v*_*z*_)^*t*^ and the pressure *P* read

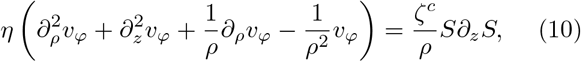

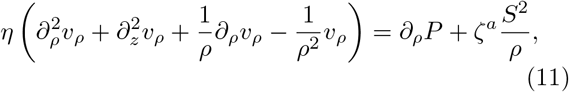

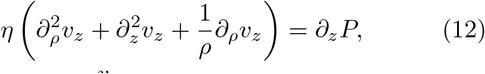

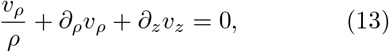

In order to numerically solve the set of equations, we assumed a cell shape where the dorsal boundary is given by

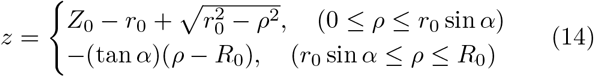

and the ventral boundary is specified with *z* = 0, as shown in Fig. S7. Here, *r*_0_ = (*R*_0_ sin *α* − *Z*_0_ cos *α*)*/*(1 − cos *α*) and *Z*_0_, *R*_0_ and *α* are the parameters that identify the cell shape. Also, since the actomyosin ring is located along the dorsal surface, in order to represent the localization numerically, we practically assume that *S*(*ρ, z*) is given by

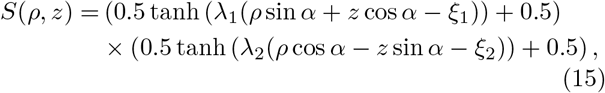

where *λ*_*i*_ and *ξ*_*i*_ (*i* = 1, 2) are parameters. An example of *S*(*ρ, z*) is shown in Fig. 6B. We numerically solved the equations of motion by assuming the no-slip boundary condition for the ventral boundary, the free slip boundary condition for the dorsal surfaces, and the vanishing flow velocity for *v*_*ρ*_ and *v*_*φ*_ and the continuity for *v*_*z*_ at the cell center *ρ* = 0. We do not include any organelles such as a nucleus in the model for simplicity. The equations were solved numerically with a finite element method using software FreeFEM++ [50]. We used the following parameter values: *η* = 1, *ζ*^*c*^ = 0.4, *ζ*^*a*^ = 1.0, *Z*_0_ = 5, *R*_0_ = 25, *α* = 15°, *λ*_1_ = 5, *λ*_2_ = 0.5, *ξ*_1_ = *R*_0_ sin *α* − 0.5, *ξ*_2_ = 11.5.

### Quantification of dorsal and ventral actomyosin

Fluorescent signals of anti-myosin IIA and phalloidin were obtained by LSM880 (Zeiss) with Airyscan and processed by ImageJ Fiji. The obtained images were resliced and ten *x-z* slices containing the cell center were processed with mean intensity projection. The dorsal and ventral surfaces were manually traced with ten-pixels-width and the average signal intensities in the traced regions were quantified. The cell edge regions of overlapping dorsal and ventral traces were annotated as “peripheral region” and excluded from the quantification.

## References

[1] H. Hamada and P. P. L. Tam, F1000prime reports 6, 110 (2014).

[2] M. Blum and T. Ott, Current Biology 28, R301 (2018).

[3] S. R. Naganathan, S. Fürthauer, M. Nishikawa, F. Jülicher, and S. W. Grill, eLife 3, e04165 (2014).

[4] L. G. Pimpale, T. C. Middelkoop, A. Mietke, and S. W. Grill, eLife 9, e54930 (2020).

[5] K. Sugioka and B. Bowerman, Developmental Cell 46, 257 (2018).

[6] Y. Shibazaki, M. Shimizu, and R. Kuroda, Current Biology 14, 1462 (2004).

[7] A. Davison, G. S. McDowell, J. M. Holden, H. F. Johnson, G. D. Koutsovoulos, M. M. Liu, P. Hulpiau, F. V. Roy, C. M. Wade, R. Banerjee, F. Yang, S. Chiba, J. W. Davey, D. J. Jackson, M. Levin, and M. L. Blaxter, Current Biology 26, 654 (2016).

[8] M. Abe and R. Kuroda, Development 146, dev175976 (2019).

[9] S. Hozumi, R. Maeda, K. Taniguchi, M. Kanai, S. Shirakabe, T. Sasamura, P. Spéder, S. Noselli, T. Aigaki, R. Murakami, and K. Matsuno, Nature 440, 798 (2006).

[10] K. Taniguchi, R. Maeda, T. Ando, T. Okumura, N. Nakazawa, R. Hatori, M. Nakamura, S. Hozumi, H. Fujiwara, and K. Matsuno, Science 333, 339 (2011), 10.1126/science.1200940.

[11] R. Hatori, T. Ando, T. Sasamura, N. Nakazawa, M. Nakamura, K. Taniguchi, S. Hozumi, J. Kikuta, M. Ishii, and K. Matsuno, Mechanisms of development 133, 146 (2014-08).

[12] K. Sato, T. Hiraiwa, E. Maekawa, A. Isomura, T. Shibata, and E. Kuranaga, Nature Communications 6, 10074 (2015).

[13] P. Ray, A. S. Chin, K. E. Worley, J. Fan, G. Kaur, M. Wu, and L. Q. Wan, Proceedings of the National Academy of Sciences 115, E11568 (2018).

[14] T. Ishibashi, R. Hatori, R. Maeda, M. Nakamura, T. Taguchi, Y. Matsuyama, and K. Matsuno, Genes to Cells 24, 214 (2019).

[15] A. Tamada, S. Kawase, S. Kawase, F. Murakami, F. Murakami, H. Kamiguchi, and H. Kamiguchi, The Journal of Cell Biology 188, 429 (2010).

[16] H. Yamanaka and S. Kondo, Genes Cells 20, 29 (2015-01).

[17] Y. H. Tee, T. Shemesh, V. Thiagarajan, R. F. Hariadi, K. L. Anderson, C. Page, N. Volkmann, D. Hanein, S. Sivaramakrishnan, M. M. Kozlov, and A. D. Bershadsky, Nature Cell Biology 17, 445 (2015-04).

[18] Y. H. Tee, W. J. Goh, X. Yong, H. T. Ong, J. Hu, I. Y. Y. Tay, S. Shi, S. Jalal, S. F. H. Barnett, P. Kanchanawong, W. Huang, J. Yan, Y. A. B. Lim, V. Thiagarajan, A. Mogilner, and A. D. Bershadsky, Nature Communications 14, 776 (2023).

[19] A. S. Chin, K. E. Worley, P. Ray, G. Kaur, J. Fan, and L. Q. Wan, Proceedings of the National Academy of Sciences 115, 12188 (2018).

[20] T.-H. Chen, J. J. Hsu, X. Zhao, C. Guo, M. N. Wong, Y. Huang, Z. Li, A. Garfinkel, C.-M. Ho, Y. Tintut, and L. L. Demer, Circulation Research 110, 551 (2012).

[21] K. Sato, T. Hiraiwa, and T. Shibata, Physical Review Letters 115, 188102 (2015).

[22] T. Yamamoto, T. Hiraiwa, and T. Shibata, Physical Review Research 2, 043326 (2020).

[23] N. A. Brown and L. Wolpert, Development 109, 1 (1990).

[24] J. Xu, A. V. Keymeulen, N. M. Wakida, P. Carlton, M. W. Berns, and H. R. Bourne, Proceedings of the National Academy of Sciences 104, 9296 (2007).

[25] T. Nishizaka, T. Yagi, Y. Tanaka, and S. Ishiwata, Nature 361, 269 (1993).

[26] I. Sase, H. Miyata, S. Ishiwata, and K. Kinosita, Proceedings of the National Academy of Sciences 94, 5646 (1997).

[27] M. Ozawa, S. Hiver, T. Yamamoto, T. Shibata, S. Upadhyayula, Y. Mimori-Kiyosue, and M. Takeichi, Journal of Cell Biology 219, e202006196 (2020).

[28] S. Tojkander, G. Gateva, and P. Lappalainen, Journal of Cell Science 125, 1855 (2012).

[29] A. L. Miller and W. M. Bement, Nature Cell Biology 11, 71 (2009).

[30] B. Nolen, N. Tomasevic, A. Russell, D. Pierce, Z. Jia, C. McCormick, J. Hartman, R. Sakowicz, and T. Pollard, Nature 460, 1031 (2009).

[31] R. Kuroda, K. Fujikura, M. Abe, Y. Hosoiri, S. Asakawa, M. Shimizu, S. Umeda, F. Ichikawa, and H. Takahashi, Scientific Reports 6, 34809 (2016).

[32] T. C. Middelkoop, J. Garcia-Baucells, P. Quintero-Cadena, L. G. Pimpale, S. Yazdi, P. W. Sternberg, P. Gross, and S. W. Grill, Proceedings of the National Academy of Sciences 118, e2021814118 (2021).

[33] S. A. Rizvi, E. M. Neidt, J. Cui, Z. Feiger, C. T. Skau, M. L. Gardel, S. A. Kozmin, and D. R. Kovar, Chemistry & Biology 16, 1158 (2009).

[34] Y. Nishimura, S. Shi, F. Zhang, R. Liu, Y. Takagi, A. D. Bershadsky, V. Viasnoff, and J. R. Sellers, Journal of Cell Science 134, jcs.253708 (2021).

[35] S. Fürthauer, M. Strempel, S. W. Grill, and F. Jülicher, The European Physical Journal E 35, 89 (2012-09).

[36] S. Fürthauer, M. Strempel, S. W. Grill, and F. Jülicher, Physical Review Letters 110, 048103 (2013).

[37] E. Tjhung, M. E. Cates, and D. Marenduzzo, Proceedings of the National Academy of Sciences 114, 4631 (2017), 1706.07723.

[38] N. D. Bade, R. D. Kamien, R. K. Assoian, and K. J. Stebe, Science Advances 3, e1700150 (2017).

[39] A. Kumar, A. Maitra, M. Sumit, S. Ramaswamy, and G. V. Shivashankar, Scientific Reports 4, 3781 (2014).

[40] Y. Nishimura, S. Shi, Q. Li, A. D. Bershadsky, and V. Viasnoff, Cells & Development 168, 203736 (2021).

[41] M. Tarama and T. Shibata, Physical Review Research 4, 043071 (2022), 2109.00841.

[42] Q. Ni, K. Wagh, A. Pathni, H. Ni, V. Vashisht, A. Upadhyaya, and G. A. Papoian, eLife 11, e82658 (2022).

[43] H. K. Kwong, Y. Huang, Y. Bao, M. L. Lam, and T.-H. Chen, ACS Biomaterials Science & Engineering 5, 3944 (2019).

[44] S. Nakamura, I. Grigoriev, T. Nogi, T. Hamaji, L. Cassimeris, and Y. Mimori-Kiyosue, PLoS ONE 7, e51442 (2012).

[45] J. Schindelin, I. Arganda-Carreras, E. Frise, V. Kaynig, M. Longair, T. Pietzsch, S. Preibisch, C. Rueden, S. Saalfeld, B. Schmid, J.-Y. Tinevez, D. J. White, V. Hartenstein, K. Eliceiri, P. Tomancak, and A. Cardona, Nature Methods 9, 676 (2012).

[46] C. Zhang, J. S. Kang, S. M. Asano, R. Gao, and E. S. Boyden, Current Protocols in Neuroscience 92, e96 (2020).

[47] C. E. Park, Y. Cho, I. Cho, H. Jung, B. Kim, J. H. Shin, S. Choi, S.-K. Kwon, Y. K. Hahn, and J.-B. Chang, ACS Nano 14, 14999 (2020).

[48] B.-C. Chen, W. R. Legant, K. Wang, L. Shao, D. E. Milkie, M. W. Davidson, C. Janetopoulos, X. S. Wu, J. A. Hammer, Z. Liu, B. P. English, Y. Mimori-Kiyosue, D. P. Romero, A. T. Ritter, J. Lippincott-Schwartz, L. Fritz-Laylin, R. D. Mullins, D. M. Mitchell, J. N. Bembenek, A.-C. Reymann, R. Böhme, S. W. Grill, J. T. Wang, G. Seydoux, U. S. Tulu, D. P. Kiehart, and E. Betzig, Science 346, 1257998 (2014).

[49] W. Thielicke and R. Sonntag, Journal of Open Research Software 9, 12 (2021).

[50] F. Hecht, Journal of Numerical Mathematics 20, 251 (2012).

