## Supplementary figures and images for "Epithelial cell chirality emerges through the dynamic concentric pattern of actomyosin cytoskeleton"

### Movie S1

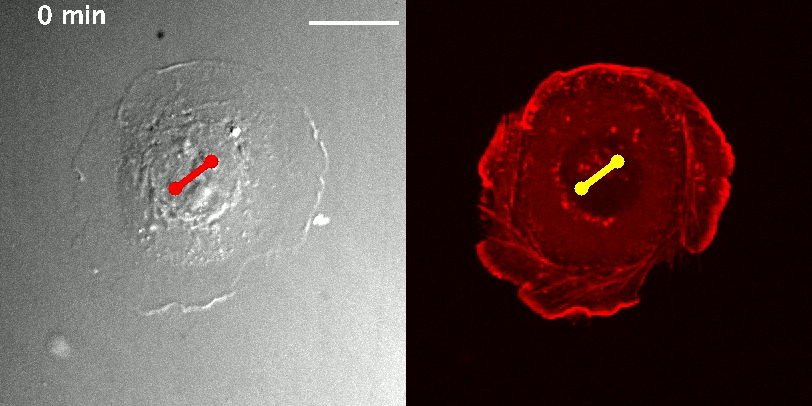

### Movie S2

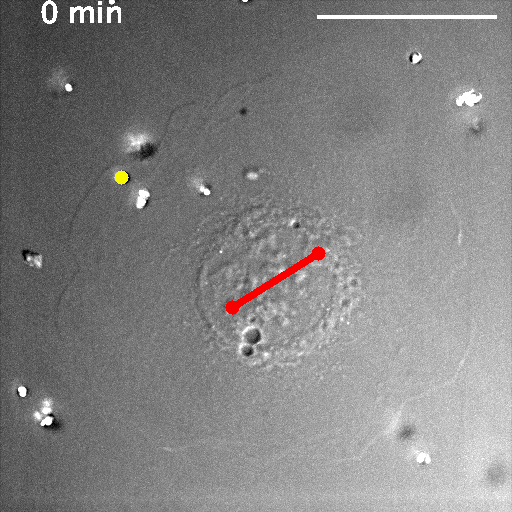

### Movie S3

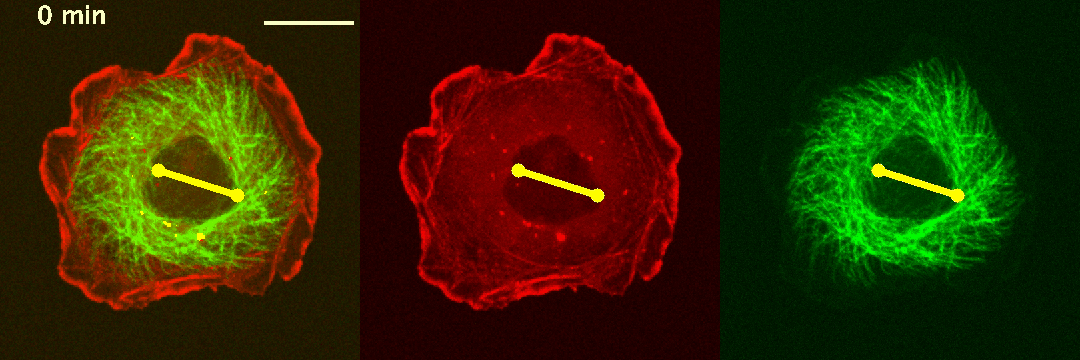

### Movie S4

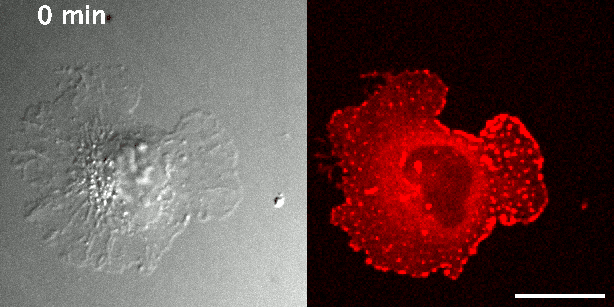

### Movie S5

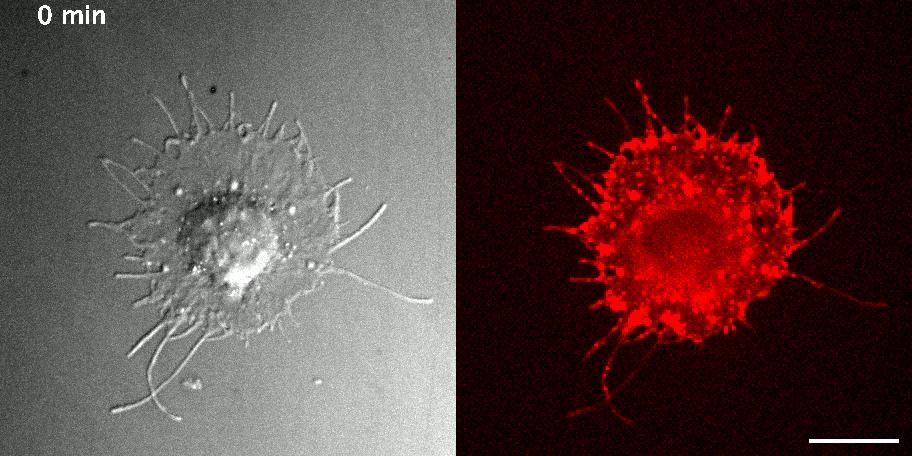

### Movie S6

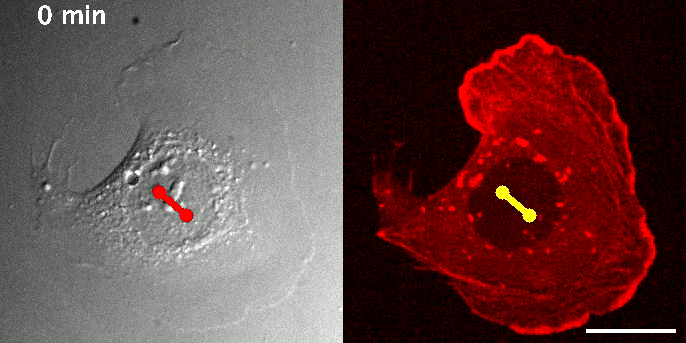

### Movie S7

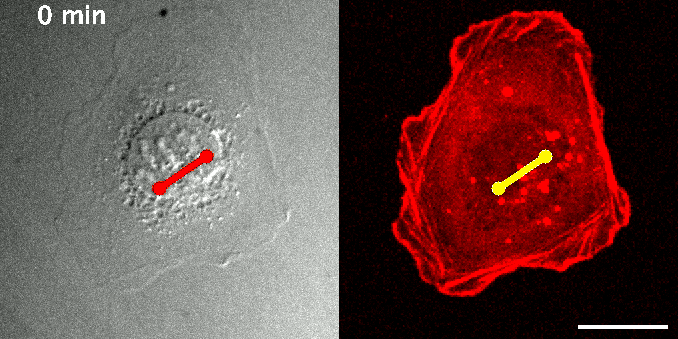

### Movie S8

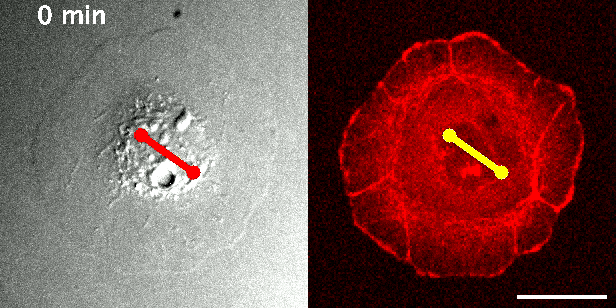

### Movie S9

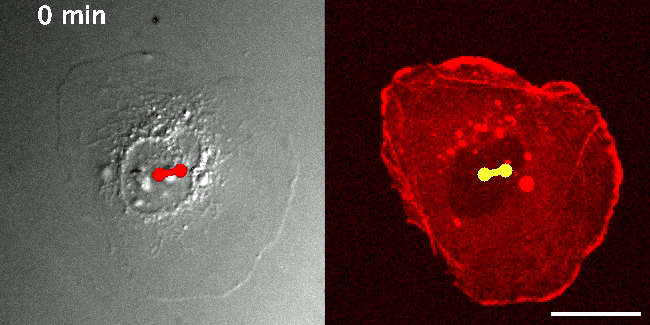

### Movie S10

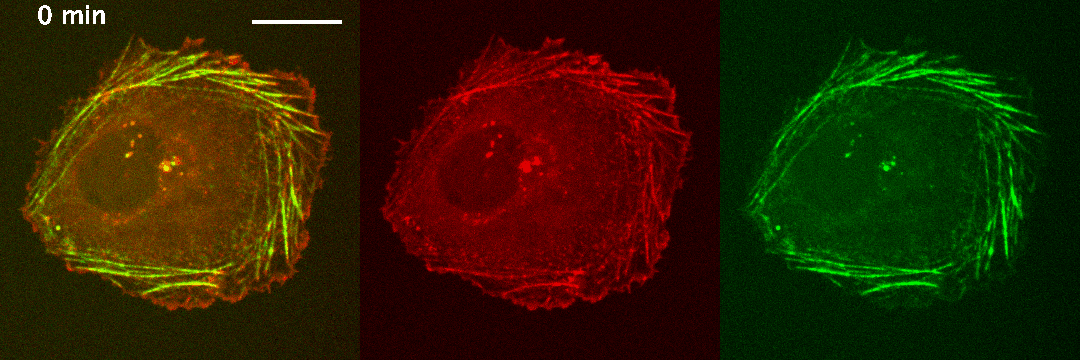

### Movie S14

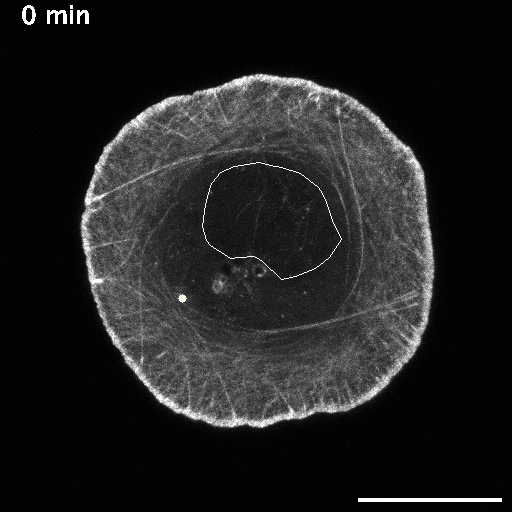

### Movie S16

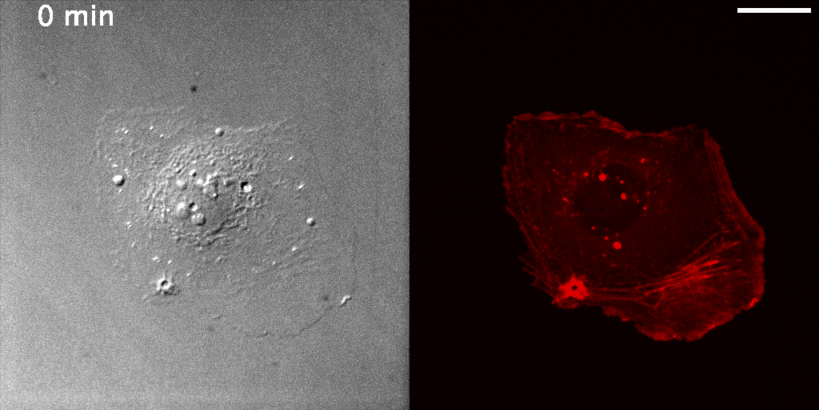
