## Supplemental Information for "Epithelial cell chirality emerges through the dynamic concentric pattern of actomyosin cytoskeleton"

**This PDF file includes:**

Figures S1 to S7
Legends for Movies S1 to S16

**Other supporting materials for this manuscript include the following:**

Movies S1 to S16

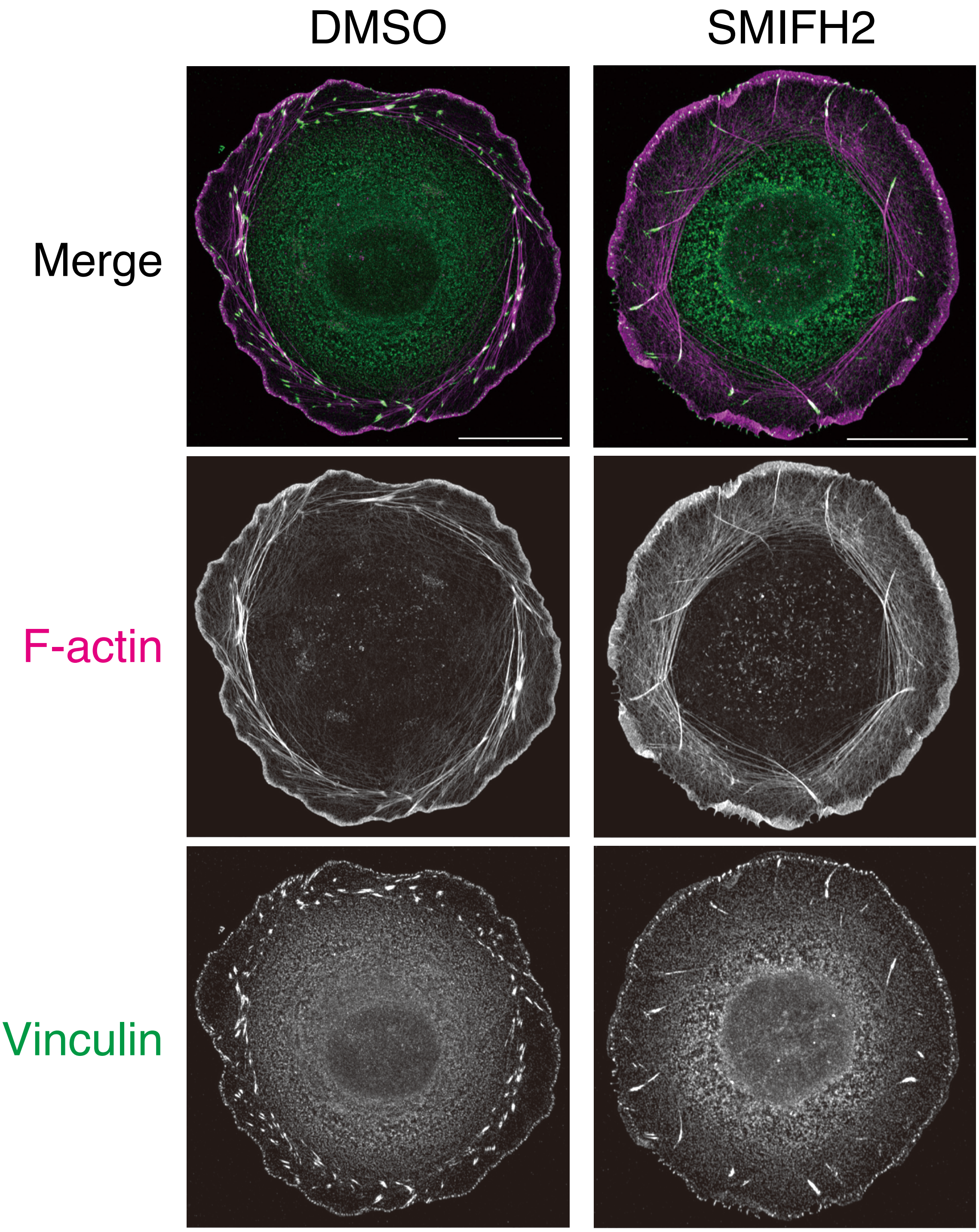


**Fig. S1**. Actin bundles in the peripheral region of cell. In the control cell (DMSO), actin bundles (phalloidin, green) in the peripheral region of cell, which are tilted to form chiral pattern, are anchored to vinculin (magenta), a focal adhesion protein, at their both ends. We call them stress fibers. In the cell treated with SMIFH2, one end of each actin bundle was anchored to vinculin, while the other ends were not anchored. Consequently, the actin bundles extend in the radial direction. We call them radial fiber.

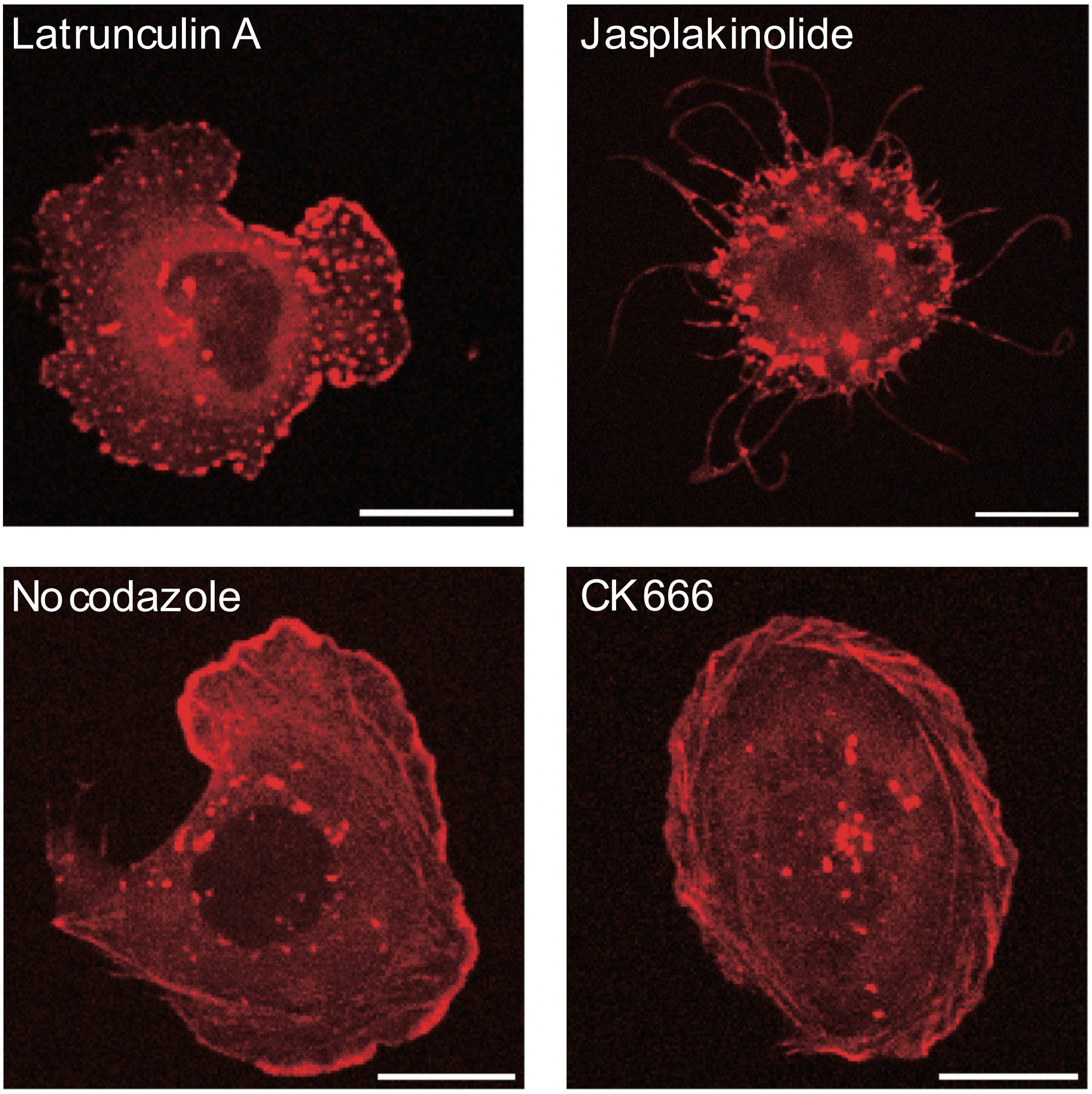


**Fig. S2**. Snapshot images from the live image of actin dynamics in cells expressing Lifeact-RFP. Cells were treated with actin polymerization inhibitor latrunculin A (2 μM), actin depolymerization inhibitor Jasplakinoilide (10 nM), microtubule inhibitor nocodazole (50 μM), or Arp2/3 inhibitor CK666 (200 μM). Scale bar: 20 μm.

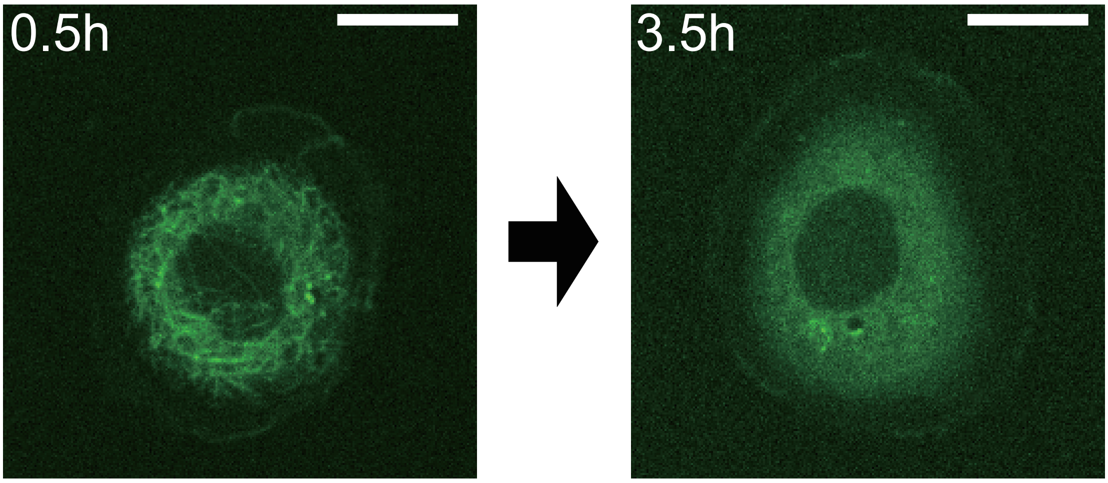


**Fig. S3.** Typical snapshots of microtubule of a cell 0.5 hour and 3.5 hour after the addition of nocodazole (50 μM). The scale bar is 20 μm.

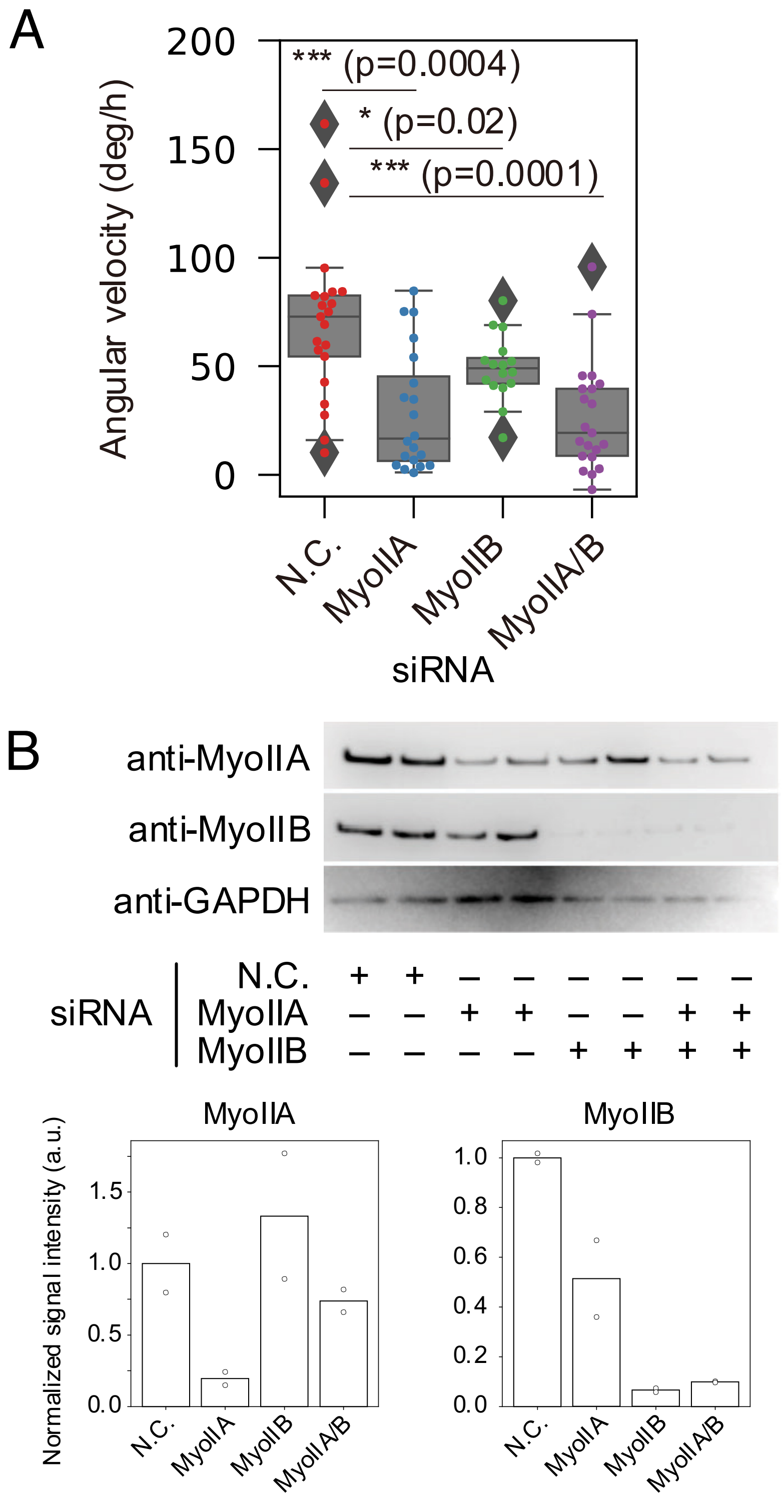


**Fig. S4.** Effect of Myosin II depletion on the nuclear rotation. (A) Angular velocity of nucleus of Caco2 cells that were treated with siRNA for myosin IIA and/or B heavy chains. The rotation of the nucleus was tracked for 10 hours following the start of the live imaging. p values were calculated using Mann-Whitney-U test ($*$ : $p < 0.05$, $**$ : $p < 0.01$, $* * *$ : $p < 0.001$). (B) Western blot showing protein levels of myosin IIA (top row) and IIB (middle row) heavy chains in Caco2 cells treated with siRNAs. GAPDH (bottom row) was used as an internal control. Quantification of fold change relative to the negative control (N.C.), normalized to GAPDH protein level, is shown in bar graphs (bottom panel).

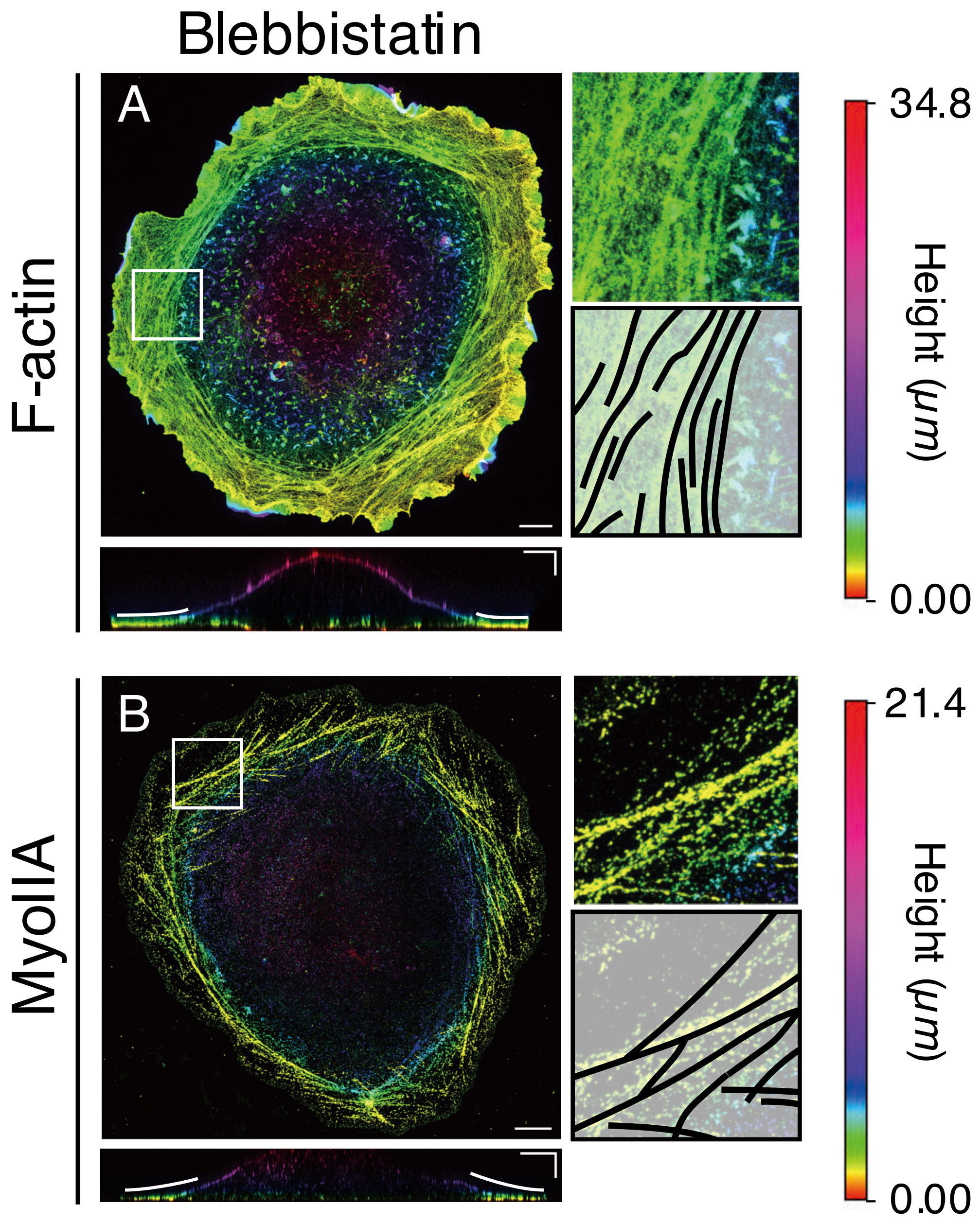


**Fig. S5.** ExM imaging of actin filaments and myosin II in cells treated with blebbistatin. Maximum intensity projection (MIP) images of F-actin (A) and myosin IIA (B) in blebbistatin-treated cells. The color indicates the height along the *z*-axis, where the height was measured after sample was swollen (color bar, the most right). Magnified views of the white boxes are shown in the right top panels, and corresponding outlines of F-actin are shown in the right bottom panels, where the bold and dotted lines indicate thick and thin fibers, respectively. The vertical section (*xz*) images are shown in the bottom panels, where the bold and dotted lines indicate peripheral and dorsal inner region, respectively. Scale bars: 20 μm (horizontal), 10 μm (vertical).

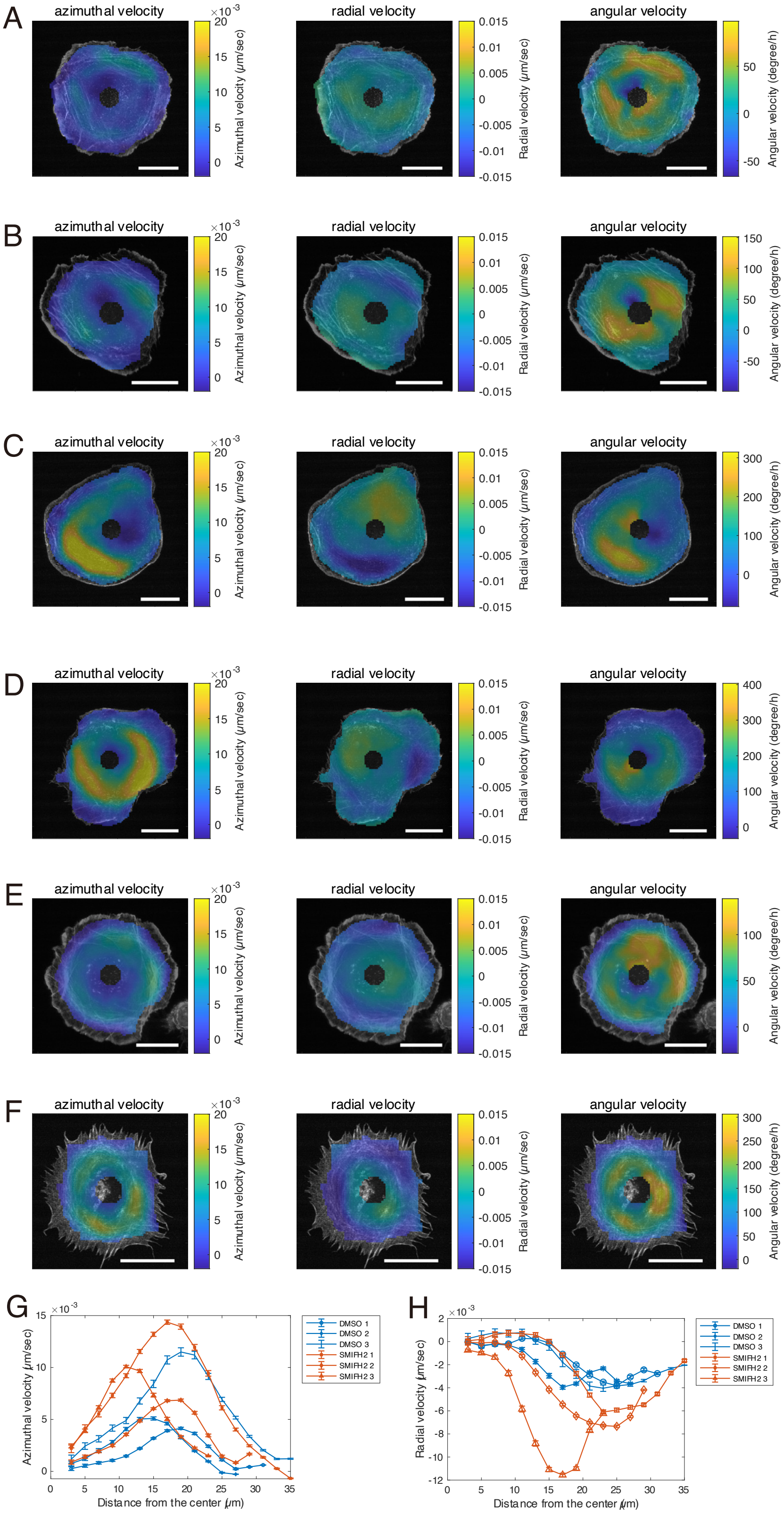


**Fig. S6.** PIV analysis of cell treated with DMSO and SMIFH2. (A-F) Spatial distribution of azimuthal velocity (left), radial velocity (middle) and angular velocity (right) for cells treated with DMSO (control) (A-C) and with SMIFH2 (D-F). (G) Angular averaged azimuthal velocity plotted against the distance from the cell center. (H) Angular averaged radial velocity plotted against the distance from the cell center. Scalebar: 20 μm. Positive azimuthal and angular velocities indicate clockwise rotation.

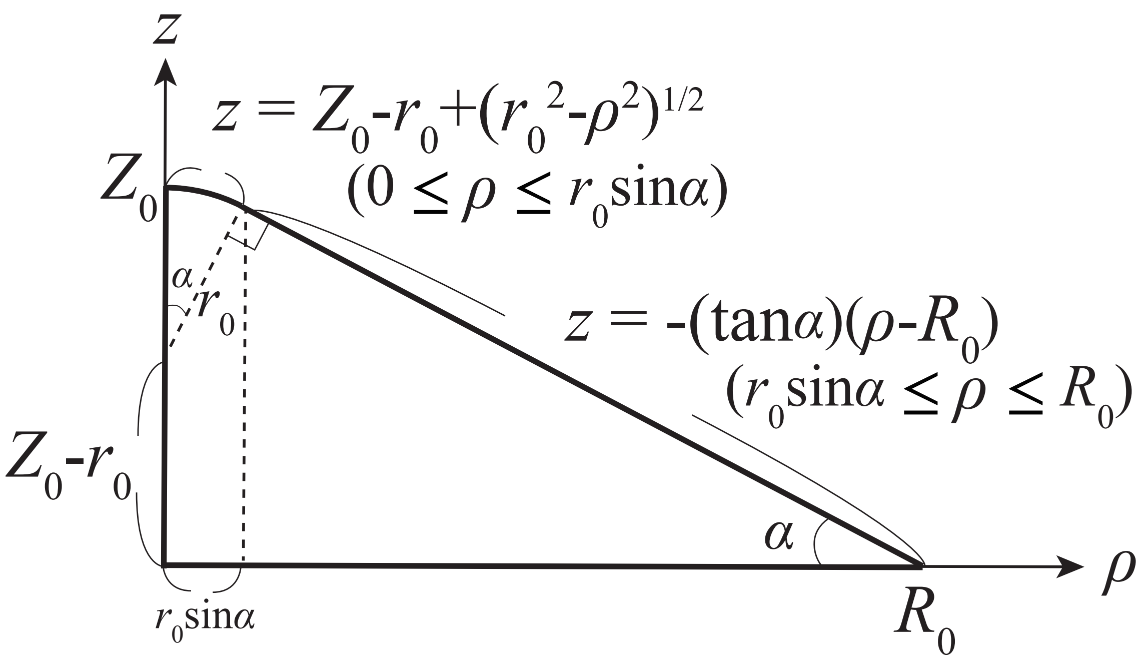


**Fig. S7.** Cell shape used in the numerical simulation. Here, cell shape is assumed to be axisymmetric with respect to the cell center. $\boldsymbol{r_0} = (\boldsymbol{R_0}\sin\boldsymbol{\alpha} - \boldsymbol{Z_0}\cos\boldsymbol{\alpha})/(\boldsymbol{1} - \cos\boldsymbol{\alpha})$ and $Z_0$, $R_0$ and $\alpha$ are the parameters to determine the cell shape.

**Movie S1 (separate file)**. DIC and fluorescent images of Caco-2 expressing Lifeact-RFP**.** Scale bar: 20 µm**.**

**Movie S2 (separate file).** DIC images of Caco-2 with beads attached to the dorsal membrane. Scale bar: 40 µm.

**Movie S3 (separate file).** Fluorescent images of Caco-2 expressing Lifeact-RFP and EMTB-3XGFP. Scale bar: 20 µm.

**Movie S4 (separate file).** DIC and fluorescent images of Caco-2 expressing Lifeact-RFP treated with latrunculin A. Scale bar: 20 µm.

**Movie S5 (separate file).** DIC and fluorescent images of Caco-2 expressing Lifeact-RFP treated with jasplakinolide. Scale bar: 20 µm.

**Movie S6 (separate file).** DIC and fluorescent images of Caco-2 expressing Lifeact-RFP treated with nocodazole. Scale bar: 20 µm.

**Movie S7 (separate file).** DIC and fluorescent images of Caco-2 expressing Lifeact-RFP treated with CK666. Scale bar: 20 µm.

**Movie S8 (separate file).** DIC and fluorescent images of Caco-2 expressing Lifeact-RFP treated with SMIFH2. Scale bar: 20 µm.

**Movie S9 (separate file).** DIC and fluorescent images of Caco-2 expressing Lifeact-RFP treated with blebbistatin. Scale bar: 20 µm.

**Movie S10 (separate file).** Fluorescent images of Caco-2 expressing Lifeact-RFP treated and MRLC-EGFP. Scale bar: 20 µm

**Movie S11 (separate file).** Fluorescent images of Caco-2 expressing Lifeact-mEmelard taken by LLSM at three different z-planes (z=0, 0.5, 1.0 µm).

**Movie S12 (separate file).** PIV analysis performed at each time step on the cell shown in Movie S11. Angular, azimuthal, and radial velocities around the cell center are indicated by color map.

**Movie S13 (separate file).** MIP Fluorescent images of Caco-2 expressing Lifeact-mEmelard treated with SMIFH2 taken by LLSM. Scale bar: 10 µm

**Movie S14 (separate file).** MIP Fluorescent images of Caco-2 expressing Lifeact-mEmelard treated with SMIFH2 taken by LSM880, Zeiss. Fluorescent debris and nucleus are marked by yellow and white line.

**Movie S15 (separate file).** MIP Fluorescent images of Caco-2 expressing Lifeact-mEmelard treated with SMIFH2 taken by LLSM. Two fluorescent debris are marked by red and yellow circles. Scale bar: 10 µm

**Movie S16 (separate file).** DIC and fluorescent images of Caco-2 expressing Lifeact-RFP treated with CN03. Scale bar: 20 µm.
